# Functional contribution of astrocytic Kir4.1 channels to spasticity after spinal cord injury

**DOI:** 10.1101/2024.10.11.617793

**Authors:** Tony Barbay, Emilie Pecchi, Jose Jorge Ramirez Franco, Anton Ivanov, Frédéric Brocard, Nathalie Rouach, Rémi Bos

## Abstract

Spasticity, a prevalent motor issue characterized by network hyperexcitability, causes pain and discomfort, with existing treatments offering limited relief. While past research has focused on neuronal factors, the role of astrocytes in spasticity has been overlooked. This study explores the potential of restoring astrocytic potassium (K^+^) uptake to reduce spasticity following SCI. Astrocytes buffer extracellular K^+^ via Kir4.1 channels, preventing neuronal hyperexcitability. Following spinal cord injury (SCI), Kir4.1 levels decrease at the injury site, though the consequences and mechanisms of this reduction within the motor output area have not been investigated. Utilizing advanced techniques, we demonstrate that lumbar astrocytes in a juvenile thoracic SCI mouse model switch to reactive phenotype, displaying morpho-functional and pro-inflammatory changes. These astrocytes also experience NBCe1-mediated intracellular acidosis, leading to Kir4.1 dysfunction and impaired K^+^ uptake. Enhancing Kir4.1 function reduces spasticity in SCI mice, revealing new therapeutic targets for neurological diseases associated with neuronal hyperexcitability.

**Highlights:**

- Lumbar astrocytes adopt a reactive phenotype following a thoracic SCI
- NBCe1-mediated acidosis in astrocytes disrupts Kir4.1 function post-SCI.
- Impaired K+ uptake leads to motoneuron hyperexcitability post-SCI.
- Enhanced astroglial Kir4.1 function reduces spastic-like symptoms in SCI mice.

## Introduction

Traumatic spinal cord injury (SCI) is characterized by an acute mechanical injury, which drastically interrupts the sensory and motor tracts, leading to a wide spectrum of chronic dysfunctions, in particular spasticity affecting ∼75% of the SCI patients^1^. Spasticity is characterized by a velocity-dependant increase in the tonic stretch reflex and spasms. It impairs the life quality of SCI patients by decreasing mobility and causing pain, sleep disturbances and depression^2^. Therapeutic approaches for spasticity are currently giving unsatisfactory results. Neurorehabilitation moderately attenuates the spastic symptoms^3^, this is why the main clinical strategy is focused on pharmacology. However, anti-spastic medications display limited benefits relative to numerous side effects. A better understanding of the pathophysiological mechanisms of spasticity is needed to propose a more appropriate therapeutic strategy.

Previous works including our own, revealed that spasticity after SCI is associated with a neuronal dysfunction resulting from an excitatory/inhibitory imbalance of both (i) pre-motor neurons^4–6^ and (ii) motoneurons (MNs) innervating the hindlimb muscles^7–13^. A switch in neuronal excitability occurs at lumbar levels where MNs switch from hypoexcitability during the initial ‘spinal shock’ to hyperexcitability a few weeks or months after SCI in adults^14^. In our newborn SCI model, we showed that neuronal hyperexcitability occurs within few days post-SCI after an initial short-term hypoexcitability^10,12^. In this model, neuronal hyperexcitability has been associated with an increase in persistent sodium induced by a calpain-driven cleavage of voltage-dependent sodium channels Nav1.6^11^. Besides this increase in neuronal excitability, we previously observed a neuronal disinhibition in MNs resulting from a decrease of KCC2 function, a cation-chloride cotransporter involved in the intracellular chloride homeostasis^8,10^. When KCC2 is reduced, the inhibitory neurotransmistters GABA and glycine contribute to hyperexcitability by evoking depolarization instead of hyperpolarization^10^. So far, most of the mechanisms proposed to explain spasticity in adult or juvenile SCI models focused on intrinsic neuronal elements. To determine the role of astrocytes, a subtype of glial cells^15^, in modulating the excitability of large-size MNs that innervate the fast-fatigable muscles, we chose a juvenile SCI model over an adult one. Indeed, recording these large-size MNs in adult lumbar spinal cord slices is extremely challenging largely due to (i) their extensive dendritic arborization^16^ and (ii) their high metabolic demands, which makes them vulnerable to ischemia^17^, a common issue during *ex vivo* slicing. To overcome these limitations, we and others, conduct studies on spinal lumbar cord slices prepared from animals within the first 2 weeks after birth ^11,12,18–20^.

In response to neuronal activity, astrocytes prevent neuronal hyperexcitability by buffering the increase in extracellular potassium concentration ([K^+^]_o_) mainly through the inwardly rectifying K^+^ channel, Kir4.1^21,22^. Recent evidences also indicate that astrocytes switching to a reactive phenotype contribute to the pathophysiology of neurodegenerative diseases and traumatic CNS injury^23^. Reactive astrocytes are defined as astrocytes that undergo morphological, molecular, and functional changes in response to pathological situations in surrounding tissue^23,24^. Among the molecular changes, a decrease in the expression of Kir4.1 channels has been observed in various neurological disorders^25^, including SCI in the area of the injury site^26^. However, the functional consequences of such a decrease within the motor output region (specifically the lumbar region where MNs are located) have not been investigated, especially in the chronic phase when spasticity emerges. Understanding the glial pathophysiological processes underlying this neuronal hyperexcitability might offer new therapeutic strategy for spasticity.

Here we investigated whether restoring astrocyte K^+^ uptake alleviates spasticity after SCI. We demonstrate that, following thoracic SCI, lumbar astrocytes switch to a reactive phenotype exhibiting morpho-functional changes and inflammatory markers. These astrocytes also display prominent NBCe1-mediated intracellular acidosis leading to dysfunction of the inwardly-rectifying Kir4.1 channels, which ultimately results in impaired K^+^ uptake, and MN hyperexcitability. Importantly, we demonstrate that enhancing Kir4.1 function alleviates spastic-like symptoms in SCI mice.

## Results

### Morphological changes of astrocytes and inflammatory marker expression in lumbar region after SCI

To determine whether astrocytic reactivity occurs in the lumbar spinal cord during the onset of spasticity following thoracic SCI, we performed immunohistofluorescence against GFAP in Aldh1L1-eGFP mice^27^, in which astrocytes express GFP (Figure 1A). This revealed significant astrocytic hypertrophy and a widespread increase in GFAP fluorescence intensity in the gray matter of the lumbar ventral horn in SCI mice (Figures 1A-1B, Figure S1A-B). Using Sholl analysis based on GFAP labelling, we showed that lumbar GFP+ astrocytes from SCI mice display longer and more branched processes, characterized by an increased ramification index (Figure 1C), and occupy a larger space than their control counterparts (Figure 1D). These results suggest the emergence of astrocyte reactivity following SCI. Given that GFAP expression and morphological changes are not conclusive indicators of reactive astrocytes^23^, we focused on additional pro-inflammatory biomarkers, namely the complement component 3d (C3d) and the transcription factor Signal Transducer and Activator of Transcrption 3 (STAT-3), both well-documented for their elevated levels in reactive astrocytes within cortical regions impacted by neurological disorders^28–30^. We found a notable rise in the proportion of Aldh1L1(+) astrocytes co-expressing the pro-inflammatory marker C3d, as shown by immunohistochemistry (Figures S1C-D). Active phosphorylated STAT3 could not be detected by immunohistochemestry in our SCI model. We used STAT3 nuclear localization as an indicator of pathway activation^29,30^, and observed a significant increase of astrocytic nuclear STAT3 translocation in SCI mice compared to controls (Figures S1E-F). Collectively, these findings reveal that thoracic SCI triggers morphological changes in astrocytes and inflammatory marker upregulation in the vicinity of lumbar ventro-lateral α-MNs.

**Figure 1.**
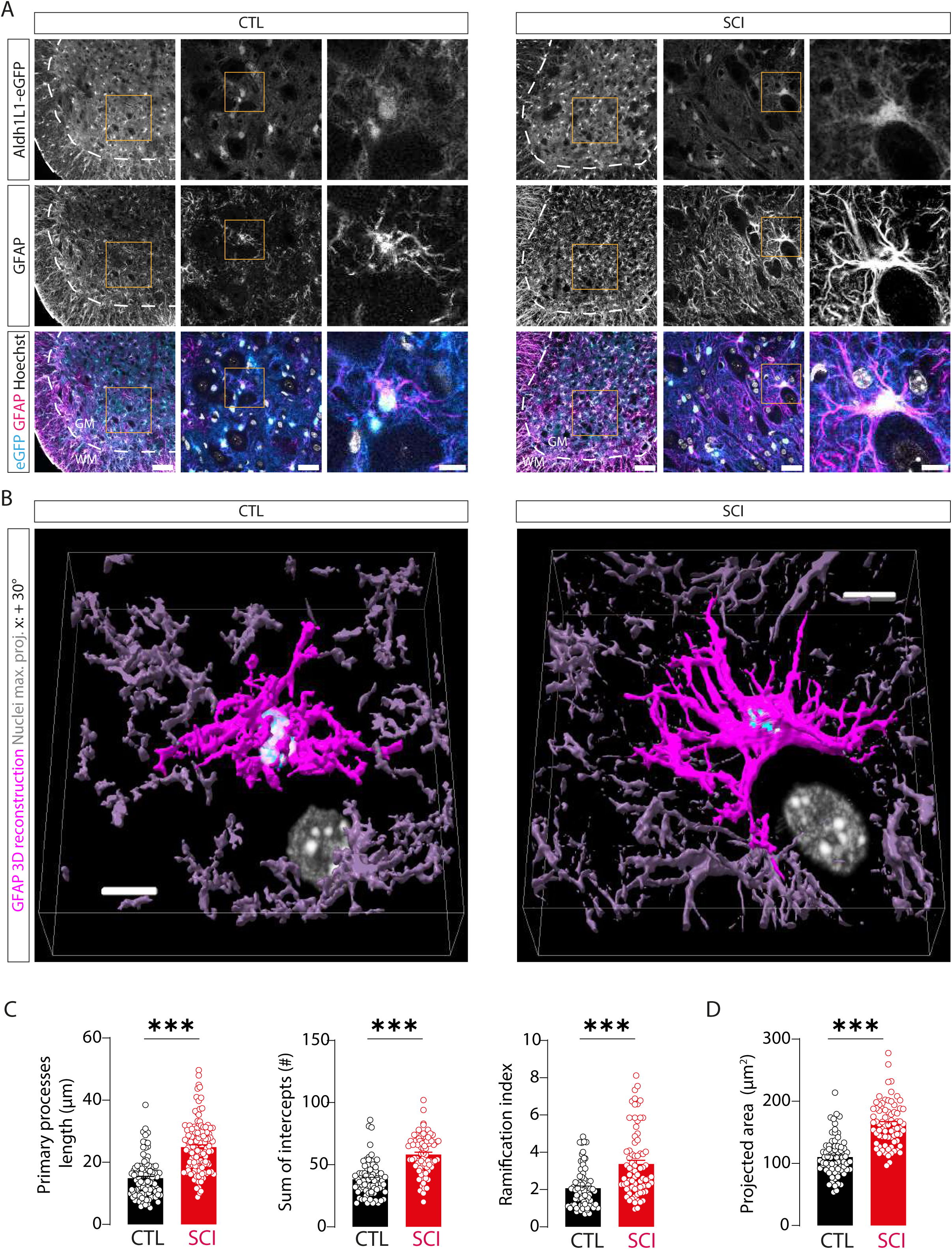
Lumbar astrocytes display morphological changes nearby motoneurons (MNs) after thoracic spinal cord injury (SCI). **(A)** Confocal images showing endogenous GFP (top) and immunofluorescent staining with GFAP (middle), and Hoechst (bottom) in the ventro-lateral lumbar cord from control (CTL) Aldh1l1-eGFP (left) or SCI Aldh1L1-eGFP (right) mice. Images at different magnifications (left to right) depict the ventral section (scale bar: 120 µm), an intermediate view of the motoneuron (MN) pool (scale bar: 40 µm), and a high magnification of a single astrocyte near a MN (scale bar: 10 µm). **(B)** 3D reconstruction (using GFAP signal) of a single astrocyte (purple) from A (left : CTL mouse, right : SCI mouse) near MNs, showing only nuclei of the astrocyte and adjacent MNs (scale bar: 10 µm). **(C-D)** Quantification of key morphological parameters of GFAP (+) astrocytes from Aldh1L1-eGFP (+) mice, including primary process length, sum of intercepts, and ramification index (C), and the projected area from a single astrocyte (D), in control (black) and SCI (red) mice (CTL: n = 4 mice, n = 20 slices, n = 76 astrocytes; SCI: n = 4 mice, n = 20 slices, n = 76 astrocytes). Data: mean ± S.E.M. ***P < 0.001, Mann-Whitney test for E. See also Figures S1.

### Functional alterations of astrocytes surrounding MNs after SCI

To evaluate the functional deficits associated with morphological alterations in astrocytes post-SCI, we conducted patch-clamp electrophysiology on GFP(+) astrocytes in acute lumbar slices from Aldh1L1-eGFP mice (Figure 2A). Our results revealed that GFP(+) astrocytes from SCI mice exhibit altered electrophysiological properties characterized by a depolarized resting membrane potential (RMP), an increased input resistance, and a decreased conductance (Figure 2B), leading to lower macroscopic currents (Figures 2C-2E) compared to controls. In addition, to better characterize these functional alterations at a cell population level, we performed two-photon imaging of astrocytic intracellular Ca^2+^ in the lumbar ventro-lateral horn (Figure 2F). Control and SCI mice were injected at birth with adeno-associated virus serotype 5 (AAV5) designed to express GCaMP6f in all astrocytic compartments (Figure 2G). We observed an increased proportion of astrocytes displaying spontaneous Ca^2+^ transients in the processes (+22%) and soma (+32%) post-SCI (Figures 2H and 2I; Figures S2A and S2B). Further analysis revealed a significant increase in the amplitude of spontaneous Ca^2+^ transients in processes and microdomains, with no change in the soma of SCI mice (Figures 2J-K; Figures S2C and S2F-H). While the duration and frequency of the spontaneous Ca^2+^ transients in soma and processes remained unchanged post-SCI, the duration decreased and the frequency increased in microdomains (Figures 2J-K; Figures S2C and S2F-H). Overall, these findings support the view that reactive astrocytes surrounding MNs undergo specific functional alterations, characterized by altered membrane properties and increased spontaneous Ca^2+^ activity, which may contribute to the hyperexcitability of MNs following SCI.

**Figure 2.**
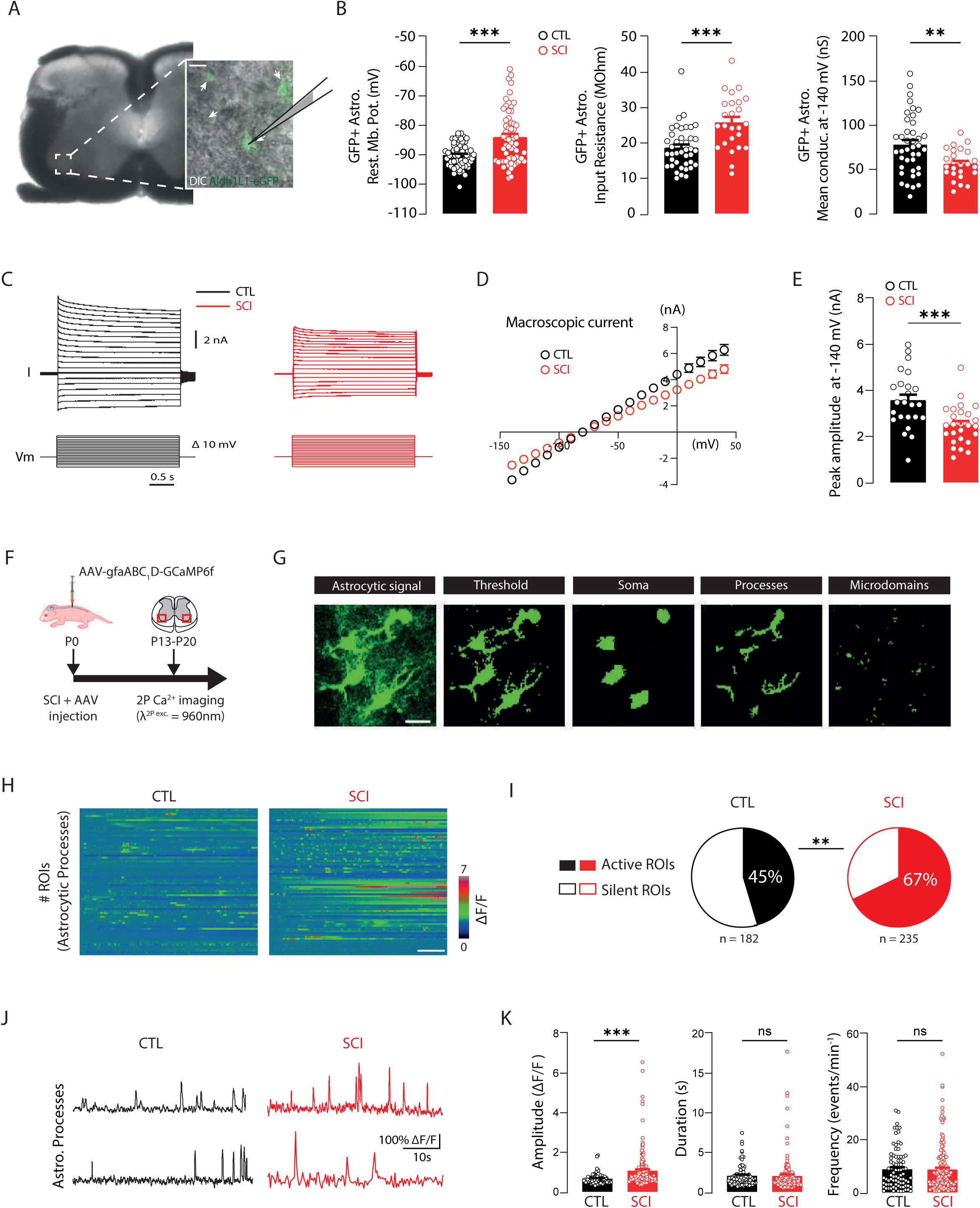
Lumbar astrocytes exhibit functional alterations after SCI. **(A)** Bright field image of a lumbar slice (L4) from an Aldh1L1-eGFP mouse under IR-DIC. Inset: High magnification of the outlined region overlaid with GFP signal. White arrows: GFP+ astrocytes. The recording pipette is black, filled in gray. Scale bar, 30 µm. **(B)** Resting membrane potential (RMP), input resistance, and mean conductance at -140mV of GFP+ astrocytes from control (CTL, black) and SCI (red) mice (CTL: n=7 mice, 61 astrocytes; SCI: n=7 mice, 60 astrocytes for RMP; CTL: n=5 mice, 40 astrocytes; SCI: n=5 mice, 25 astrocytes for resistance and conductance). **(C)** Macroscopic currents of GFP+ astrocytes clamped at RMP in response to voltage steps (-140 to +40mV) from CTL (black) and SCI (red) mice. **(D)** Quantification of macroscopic currents as a function of voltage steps. **(E)** Peak amplitude of macroscopic current at -140mV (CTL: n=5 mice, 23 astrocytes; SCI: n=5 mice, 25 astrocytes). **(F)** Experimental design for viral infection at birth followed by Ca^2+^ imaging (GCaMP6f) two to three weeks later. **(G)** Astrocyte segmentation for sub-cellular compartment analysis (scale bar: 10 µm). **(H)** Heatmaps of spontaneous astrocytic processes ROIs transients (CTL: n=182 ROIs, 7 mice; SCI: n=235 ROIs, 5 mice). Color code: fluorescence changes (ΔF/F) (scale bar: 30 s). **(I)** Percentage of active versus silent astrocytic processes (CTL: n=182 ROIs, 7 mice; SCI: n=235 ROIs, 5 mice). **(J)** Examples of spontaneous GCaMP6f Ca^2+^ transients in astrocyte processes from CTL (left) and SCI (right) mice. **(K)** Quantification of the spontaneous GCaMP6f Ca^2+^ transients amplitude, duration, and frequency in astrocytic processes (CTL: n=83 ROIs, 7 mice; SCI: n=153 ROIs, 5 mice). Data: mean ± S.E.M. ns: not significant, **P < 0.01, ***P < 0.001, Mann-Whitney test for B, E, and K. Fisher’s test for I. See also Figure S2.

### Glutamate-evoked Ca^2+^ transients are enhanced in astrocytes and MNs after SCI

Spasticity resulting from SCI is partly attributed to MNs hyperexcitability^4,8,10,11^. Given that reactive astrocytes are known to modulate neuronal excitability^31,32^, we investigated the SCI-induced changes in astrocyte-neuron signaling. To address this, we conducted simultaneous two-photon Ca^2+^ imaging of astrocytes and neurons in the ventro-lateral lumbar region. At birth, wild-type mice were injected with AAV5 to express GCaMP6f in astrocytes (Figure 3A) and AAV9 to express jRGECO1a in neurons (Figure 3B). Glutamate was bath applied to trigger widespread increases in neuronal activity. Following SCI, we observed significantly elevated Ca^2+^ transients in astrocytic processes (+214%), soma (+166%) and microdomains (+139%) in response to glutamate exposure, compared to control conditions (Figures 3C-E, Figures S2D-E and S2I-J). Additionally, large neurons in the ventro-lateral region, likely α-MNs, exhibited an amplified glutamate-induced Ca^2+^ signals (+34%) (Figures 3F-3H). This pronounced Ca^2+^ activity spread progressively throughout both astrocytic and neuronal networks (Figures 3I-J). Notably, we observed that the number of co-active astrocytes is transiently increased upon glutamate exposure (+44% and +74% at 10 and 30s, respectively) in SCI mice compared to controls (Figure 3J). Altogether, these findings suggest that SCI amplifies astrocyte-neuron Ca^2+^ signaling in the lumbar motor network.

**Figure 3.**
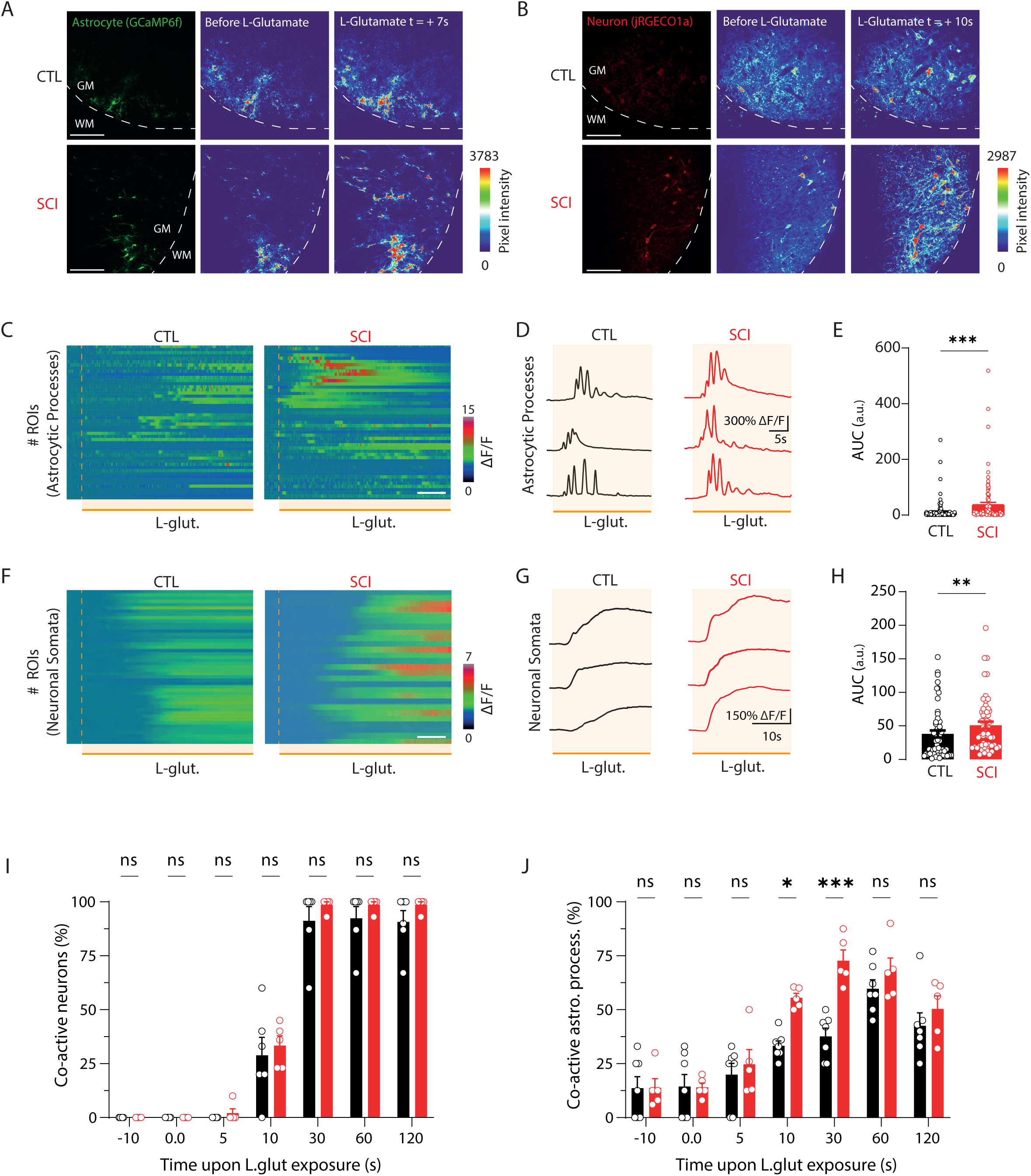
Simultaneous two-photon Ca^2+^ imaging of astrocytic and neuronal signals reveals increased glutamate-evoked Ca^2+^ transients after SCI. **(A-B)** Fluorescence images of the ventro-lateral lumbar slice showing astrocytic GCaMP6f (A) and neuronal jRGECO1a (B) Ca^2+^ signals from CTL (top left) and SCI (bottom left) mice. Pseudocolor images display Ca^2+^ transients before (middle) and after (right) L-glutamate (1mM) bath application in CTL (top) and SCI (bottom) mice. Color changes indicate pixel intensity variations. **(C)** Heatmaps of glutamate-evoked GCaMP6f Ca^2+^ transients in astrocytic processes from CTL (left) and SCI (right) mice, with color representing fluorescence changes (ΔF/F). Scale bar: 30 s. **(D)** Examples of glutamate-evoked GCaMP6f Ca^2+^ transients in astrocyte processes from CTL (left) and SCI (right) mice. **(E)** Quantification of astrocytic area under the curve (AUC) of glutamate-evoked GCaMP6f Ca^2+^ transients in CTL (black) and SCI (red) mice (CTL: n=121 ROIs, 7 mice; SCI: n=95 ROIs, 5 mice). **(F)** Heatmaps of glutamate-evoked jRGECO1a Ca^2+^ signals in neuronal soma from CTL (left) and SCI (right) mice, with color representing fluorescence changes (ΔF/F). Scale bar: 30 s. **(G)** Examples of glutamate-evoked jRGECO1a Ca^2+^ signals in neuronal soma from CTL (left) and SCI (right) mice. **(H)** Quantification of neuronal AUC of glutamate-evoked jRGECO1a Ca^2+^ transients in neuronal soma from CTL (black) and SCI (red) mice (CTL: n=54 ROIs, 6 mice; SCI: n=57 ROIs, 5 mice). **(I)** Quantification of the co-active neurons (%) in function of the time upon L-glutamate exposure (CTL: n= 6 mice; SCI: n= 5 mice). **(J)** Quantification of the co-active astrocyte processes (%) in function of the time upon L-glutamate exposure (CTL: n= 7 mice; SCI: n= 5 mice). Data: mean ± S.E.M. **P < 0.01, ***P < 0.001, Mann-Whitney test for E and H. Two-way ANOVA with Sidak’s multiple comparaison test for I and J. See also Figure S2.

### Glutamate-evoked astrocytic acidosis is enhanced after SCI

In response to increased neuronal activity, astrocytes express a high level of electrogenic sodium (Na^+^) – bicarbonate (HCO3^−^) cotransporter 1 (NBCe1, *Slc4a4*) to maintain brain extracellular pH (pHe) homeostasis^33^. NBCe1 is a high affinity carrier primarily responsible for moving HCO3^−^ across the astroglial membrane, and its activity is modulated by intracellular Ca^2+^ rise^34,35^. Given our observation of increased Ca^2+^ signals in astrocytes post-SCI, we investigated whether NBCe1 expression and function in spinal astrocytes are affected by SCI. We found a slight but significant upregulation of membrane-bound NBCe1 in astrocytes after SCI (Figures 4A and 4B). Consistent with our previous experiment, we activated the neuronal network with glutamate. Using two-photon imaging with the pH-sensitive probe BCECF during glutamate exposure, we monitored pH shifts in Td-tomato (+) astrocytes in the ventro-lateral region of lumbar slices (Figure 4C). Confirming BCECF pH-sensitivity (Figures S3A-C), astrocytes loaded with BCECF exhibited either alkalinization or acidification in response to glutamate (Figures 4D and 4E). We found a significant increase (∼20%) in acidic responses in astrocytes post-SCI compared to control (CTL) mice (Figure 4F). Detailed analysis showed a pronounced glutamate-induced functional acidification of astrocytes post-SCI, attenuated by NBCe1 blockade (Figures 4G and 4H). These data suggest that reactive astrocytes exhibit increased NBCe1 expression and undergo prominent intracellular acidosis during sustained neuronal activity post-SCI.

**Figure 4.**
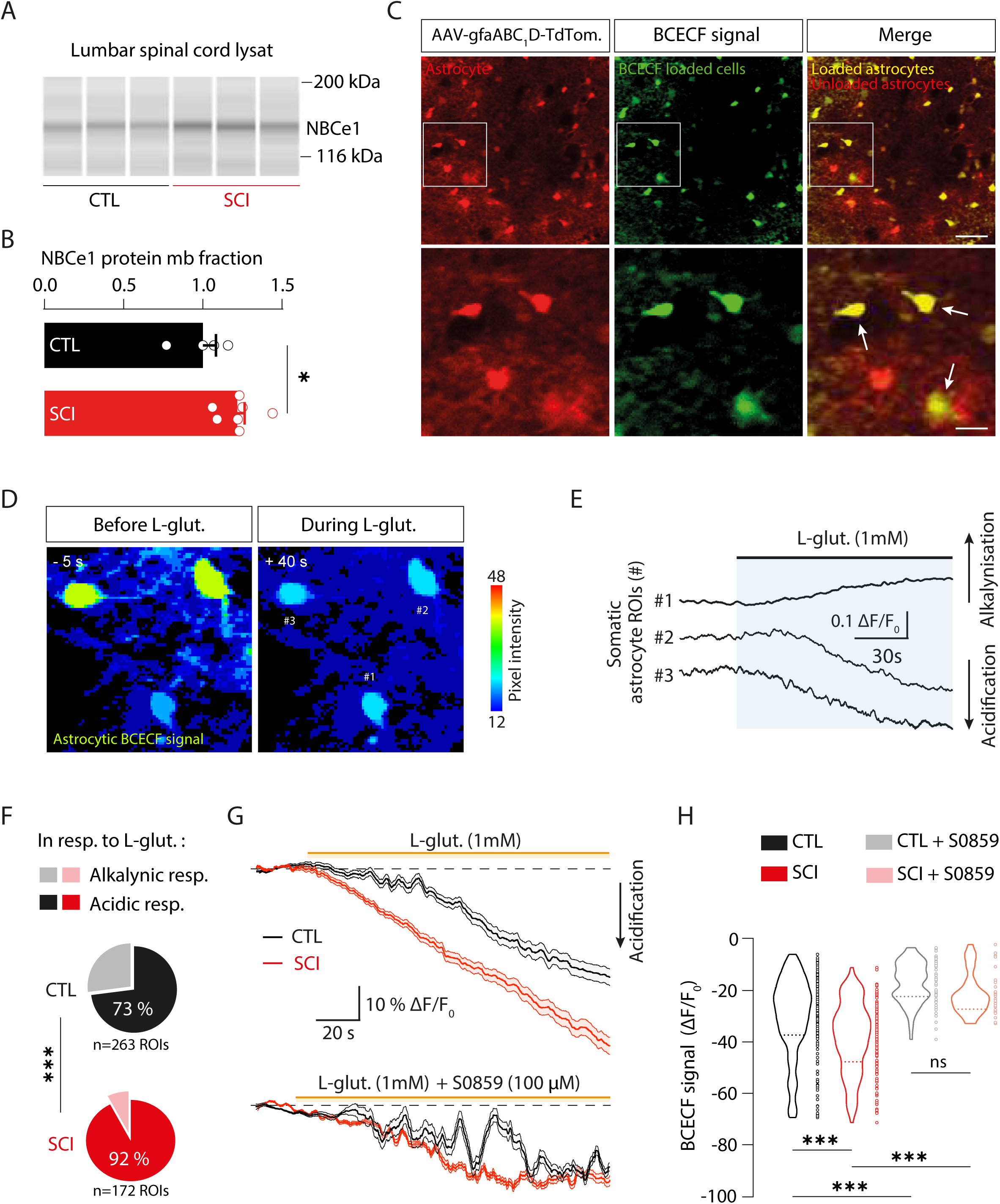
Astrocytes display increased NBCe1 levels and undergo prominent glutamate-evoked acidosis after SCI. **(A)** NBCe1 pseudo-gel images from capillary western blot of lumbar segment from CTL (left, black) and SCI (right, red) mice. **(B)** Group mean quantification of the NBCe1 band (∼140 kDa) normalized to CTL group (CTL: n=4 mice; SCI: n=7 mice). **(C)** Two-photon fluorescence images of lumbar cord showing identified astrocytes (tdTomato) and cells loaded with pH-sensitive BCECF. Scale bar: 50 µm; inset: BCECF-loaded astrocytes (white arrows). Scale bar : 15 µm. **(D)** Pseudocolor images of BCECF signals in tdTomato(+) astrocytes before (left) and after 40s of L-glutamate application (right). ROI#1 shows increased pixel intensity, while ROIs #2 and #3 show decreased signal intensity in response to L-glutamate. **(E)** BCECF signal changes in response to L-glutamate in astrocytic ROIs from the same FOV in (D), showing alkalinization (top) or acidification (bottom). **(F)** Percentage of alkalinizing and acidifying responses to glutamate in astrocytes from CTL (top, black) and SCI (bottom, red) mice (CTL: n=263 ROIs, 4 mice; SCI: n=172 ROIs, 4 mice). **(G)** (Top) Average astrocyte acidic responses during L-glutamate exposure in aCSF showing enhanced acidosis post-SCI; (Bottom) Reduced response with NBCe1 blocker, S0859 (100 µM). **(H)** Violin plots of maximum BCECF acidic changes during L-glutamate exposure in CTL (left, black) and SCI (right, red) mice without (left) or with (right) S0859 (CTL: n=158 ROIs, 4 mice; SCI: n=139 ROIs, 4 mice; with S0859, CTL: n=64 ROIs, 4 mice; SCI: n=32 ROIs, 3 mice). Data: mean ± S.E.M, hatched line in violin plots are median. ns: not significant, *P < 0.05, ***P < 0.001. Mann-Whitney test for B; Kruskal-Wallis one-way ANOVA followed by Dunn’s post-hoc test for H; Fisher’s test for F. See also Figure S3.

### NBCe1-mediated prominent acidosis in astrocytes decreases Kir4.1 function after SCI

Intracellular pH modulates various cellular functions, including the weakly inwardly rectifying K^+^ (Kir4.1) channel activity^36,37^. Given the alterated astrocytic pH regulation post-SCI, we investigated its impact on Kir4.1 functionality in reactive astrocytes. Immunohistochemistry confirmed the presence of Kir4.1 channels in astrocytes enwrapping ventro-lateral MNs in the lumbar spinal segments (L4-L5) (Figure 5A). In parallel, we assessed their pH sensitivity in our preparation. Patch-clamp recordings of GFP(+) astrocytes in the ventro-lateral lumbar slices of Aldh1L1-eGFP mice revealed a barium (Ba^2+^,100µM) -sensitive current indicative of Kir4.1 function, which was significanlty reduced under intracellular acidic conditions or post-SCI (Figures 5B-E). Interestingly, blockade of NBCe1 with S0859 (50µM) restored Kir4.1 conductance, highlighting the acidosis’ detrimental effect on Kir4.1 function (Figures 5C-E). Then, we challenged GFP(+) astrocytes with high extracellular K^+^ puff application (Figure 5F) to analyze K^+^ inward current dynamics. We first confirmed that blockade of Kir4.1 channels with Ba^2+^ (100µM) significantly decreases the astrocytic K^+^-mediated inward current (Figures S4A-C). Additionally, the application of a K^+^ puff (12 mM) induced an inward K^+^ current, which was reduced in SCI astrocytes (Figures 5G and 5H). This decrease of the pH-sensitive current was restored by blocking NBCe1. These results suggest that the NBCe1-mediated acidosis impairs K^+^ uptake by decreasing Kir4.1 function in reactive astrocytes post-SCI.

**Figure 5.**
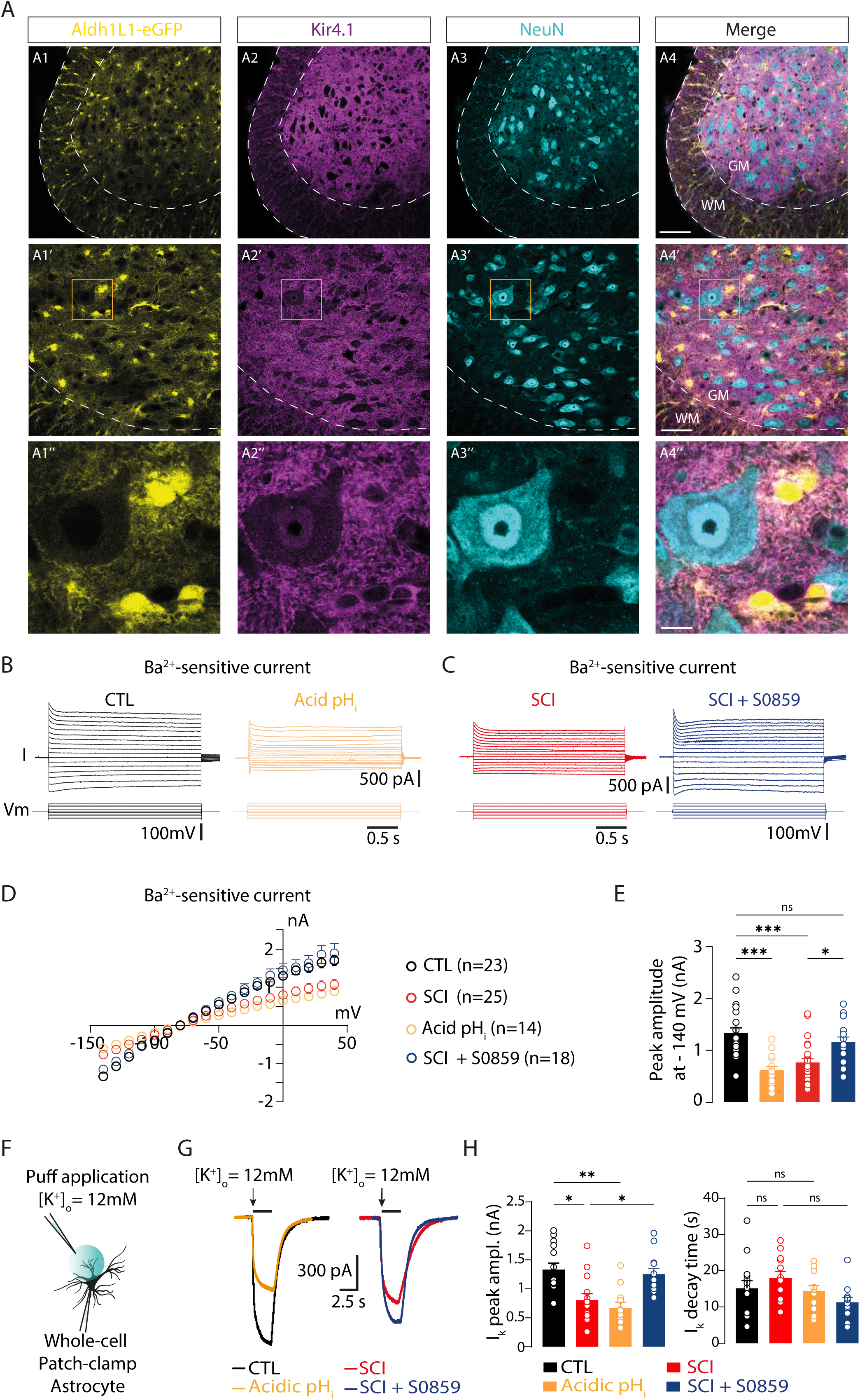
Increased NBCe1-mediated astrocytic acidosis following SCI impairs Kir4.1 function. **(A)** Confocal images showing endogenous GFP (A1), and immunofluorescent staining for Kir4.1 (A2), NeuN (A3), and merged image (A4) in the ventro-lateral lumbar cord of an Aldh1L1-eGFP mouse. Scale bar: 120 µm. Higher magnification (middle) shows astrocytes (A1’) expressing Kir4.1 (A2’) among neurons (A3’). Scale bar: 50 µm. Insets: close-up of astrocytes (A1’’) expressing Kir4.1 (A2’’) enwrapping a large ventro-lateral neuron, likely motoneurons (A3’’). Scale bar: 10 µm. GM: gray matter; WM: white matter. **(B)** Superimposed Ba^2+^-sensitive currents (top) from GFP+ astrocytes clamped at RMP in response to voltage pulses (bottom) in CTL mice with classic (pH=7.4, left) and acidic (pH=6.9, right) intracellular solution. **(C)** Superimposed Ba^2+^-sensitive currents (top) from a GFP+ astrocyte at RMP with voltage pulses (bottom) in an SCI mouse without (left) and with (right) NBCe1 blocker, S0859. **(D)** I/V plots of Ba^2+^-sensitive currents in CTL (black), CTL acidosis (yellow), SCI (red), and SCI + S0859 (blue). **(E)** Peak amplitude at -140 mV from Ba^2+^-sensitive currents (CTL: n=23 astrocytes, 5 mice; CTL pH acidosis: n=14 astrocytes, 4 mice; SCI: n=25 astrocytes, 5 mice; SCI + S0859: n=18 astrocytes, 5 mice). **(F)** Experimental design diagram for recording K^+^-inward currents in astrocytes. **(G)** Typical K^+^-inward current traces following a K^+^ puff (12mM) in astrocytes from CTL mice without (black) and with (yellow) acidic solution, and SCI mice without (red) and with (blue) S0859. (H) Peak amplitude of K^+^ puff-mediated currents (CTL: n=14 astrocytes, 5 mice; CTL pH acidosis: n=12 astrocytes, 4 mice; SCI: n=14 astrocytes, 5 mice; SCI + S0859: n=12 astrocytes, 4 mice). Data: mean ± S.E.M. ns: not significant, *P < 0.05, **P < 0.01, ***P < 0.001, Kruskal-Wallis one-way ANOVA followed by Dunn’s post-hoc test for E and H. See also Figure S4.

### Astrocytic Kir4.1 dysfunction leads to impaired K^+^ homeostasis post-SCI

MN hyperexcitability occurs within a few days post-SCI in a newborn SCI model^10,12^. It has been shown that an effective regulation of extracellular K^+^ concentration ([K^+^]_o_) is essential for limiting network hyperexcitability^38^, with astrocytes primarily maintaining K^+^ homeostasis through the activity of Kir4.1 channels^21,25^. Given that SCI induces Kir4.1 dysfunction, we investigated whether extracellular K^+^ homeostasis was impaired in lumbar slices of SCI mice. To do so, we used K^+^-sensitive microelectrodes to measure glutamate-induced changes in extracellular K^+^ concentration in control and SCI lumbar slices, specifically in the ventro-lateral region where MNs are located (Figure 6A). The [K^+^]_o_ transients evoked by glutamate puff (Figure 6B) showed a larger peak amplitude and area in SCI mice compared to controls while their rise slope and decay time were identical in both groups (Figure 6C). These findings indicate impaired glutamate-evoked K^+^ homeostasis in the lumbar cord post-SCI. This prompted us to explore the impact of increased K^+^ on the properties of large lumbar α-MNs in control mice (Figure 6D). We found that the rise in [K^+^]_o_ that we observed in SCI condition, enhanced MNs excitability by depolarizing their resting membrane potential (RMP) (Figure 6E) and decreasing their rheobase (Figures 6F), leading to a gain increase (Figure 6G). Additionally, we observed that the rise in [K^+^]_o_ decreased the peak amplitude of the slow inactivating outward K^+^ currents, primarly mediated by the Kv1.2 channels, which may contribute to the amplification of the motor output^18^. Altoghether, these findings demonstrate that SCI impairs K^+^ homeostasis in the lumbar cord, which in turn enhances MN excitability.

**Figure 6.**
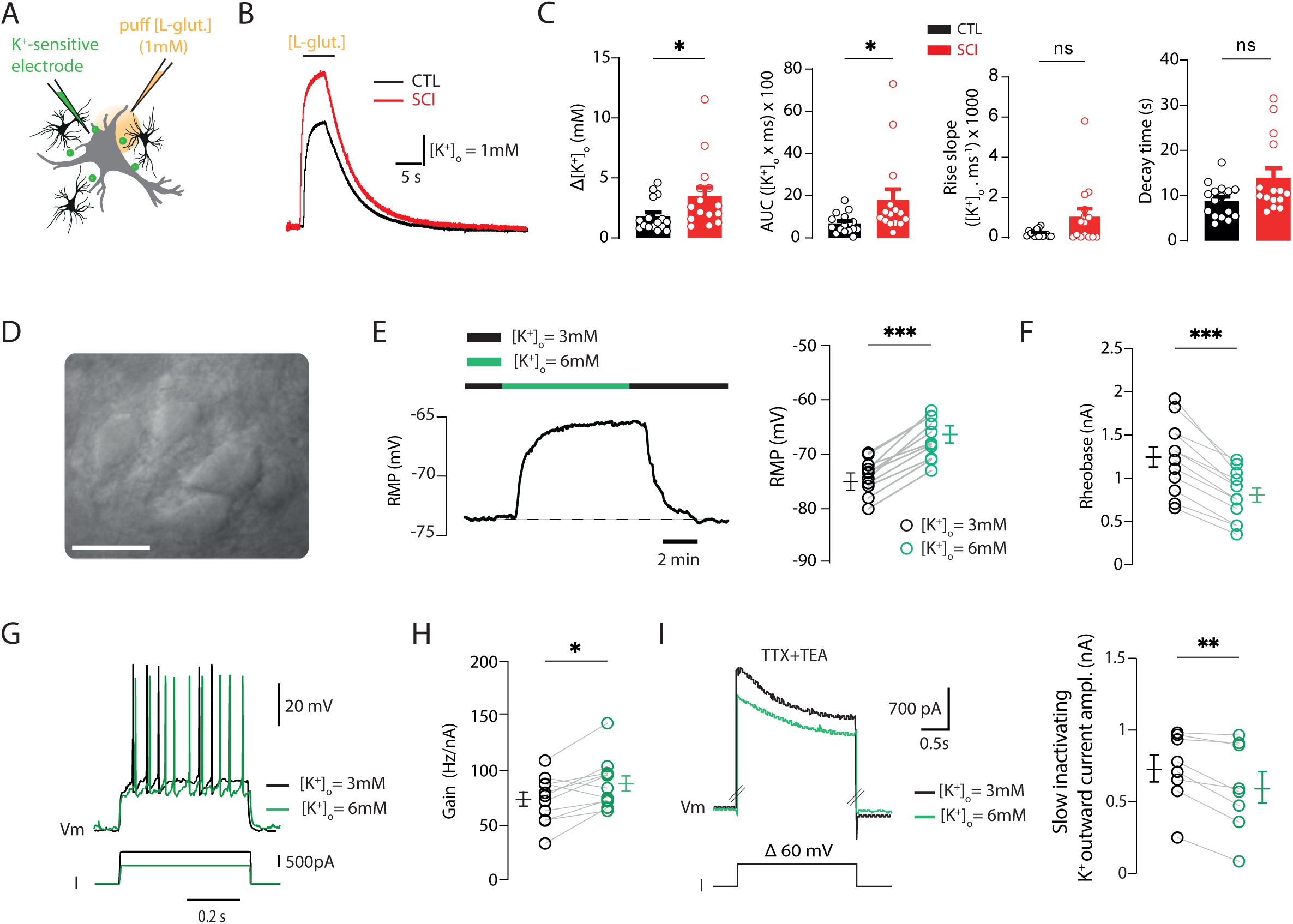
Impaired K^+^ homeostasis mediates MN hyperexcitability after SCI. **(A)** Schematic of experimental procedure. **(B)** Extracellular potassium concentration ([K^+^]_o_) increase following glutamate puff in the ventro-lateral horn of the lumbar cord from control (black) and SCI (red) mice. **(C)** Quantification of peak amplitude (left), area under the curve (AUC) (middle left), rise slope (middle right), and decay time (right) of [K^+^]_o_ variation mediated by glutamate (CTL: n=16 slices, 7 mice; SCI: n=15 slices, 5 mice). **(D)** High magnification bright field image of an acute lumbar slice from a control mouse showing a pool of motoneurons (MNs) under IR-DIC. Scale bar : 40µm. **(E)** Illustration (left) and quantification (right) of reversible MN membrane depolarization mediated by [K^+^]_o_ = 6mM (green) (n=12 neurons, 4 mice). **(F)** Quantification of rheobase before (black) and during (green) [K+]o = 6mM. **(G)** Voltage traces from MNs recorded in response to incremented current steps before (black) and during (green) bath perfusion of [K^+^]_o_ = 6mM. **(H)** Quantification of the firing gain before (black) and during (green) [K^+^]_o_ = 6mM (n=11 neurons, 4 mice). **(I)** Superimposed outward K^+^ currents in presence of TTX (1µM) and TEA (10mM) elicited by a depolarizing step (60mV) before (black) and during (green) [K^+^]_o_ = 6mM (n=8 neurons, 3 mice). Data: mean ± S.E.M. ns: not significant, *P < 0.05, **P < 0.01, ***P < 0.001, Mann-Whitney test for C, Wilcoxon test for E, F, H, and I.

### Restoration of astrocytic Kir4.1 function reduces spasticity post-SCI

To evaluate whether restoring Kir4.1 function in astrocytes could alleviate spastic-like symptoms post-SCI, we performed EMG recordings of the *gastrocnemius* muscle in wild-type SCI mice (Figure 7A), injected at birth with AAV9 carrying GFP-tagged Kir4.1 (Kir4.1-GFP) or GFP-tagged Cre (Control-GFP) under the astrocytic promoter, gfaABC1D or GFAP, respectively. First, we confirmed that Kir4.1-GFP transduction increases Kir4.1 expression in the lumbar spinal cord (Figures S5A-D), while no Kir4.1-GFP transduction occurs in the supraspinal structures involved in motor control (Figure S5E). Then, we observed a significant reduction in spontaneous muscle spasm frequency from P15 in Kir4.1-GFP-treated SCI mice, although spasm duration remained unchanged (Figures 7C ; Figures S6A and S6B). Using sensory-evoked stimulation, we found a reduced duration of triggered spasms from P10 in Kir4.1-GFP-treated SCI mice (Figures 7D and 7E). Complementary DeepLabCut analysis^39^ (Figures 7F; Figures S6C and S6D) demonstrated several additional differences between the Kir4.1-GFP-treated SCI mice and their control counterparts: (i) a decrease in spontaneous hindlimb motion frequency and acceleration from P10 and P20, respectively (Figure 7G), (ii) a significant decrease in duration of the hindlimb hyperextension from P10 in Kir4.1-GFP-treated SCI mice (Figure 7H), confirming our EMG recordings. Importantly, the tail pinch stimulation was consistent accross both conditions (Figures S6Fand S6G).and no change in ankle angle variation in response to tail pinch was observed in both conditions (Figure 7H; Figure S6E). These *in vivo* results suggest that restoring Kir4.1 function in astrocytes decreases spastic-like symptoms, suggesting Kir4.1 as a therapeutic target for neurological disorders characterized by neuronal hyperexcitability.

**Figure 7.**
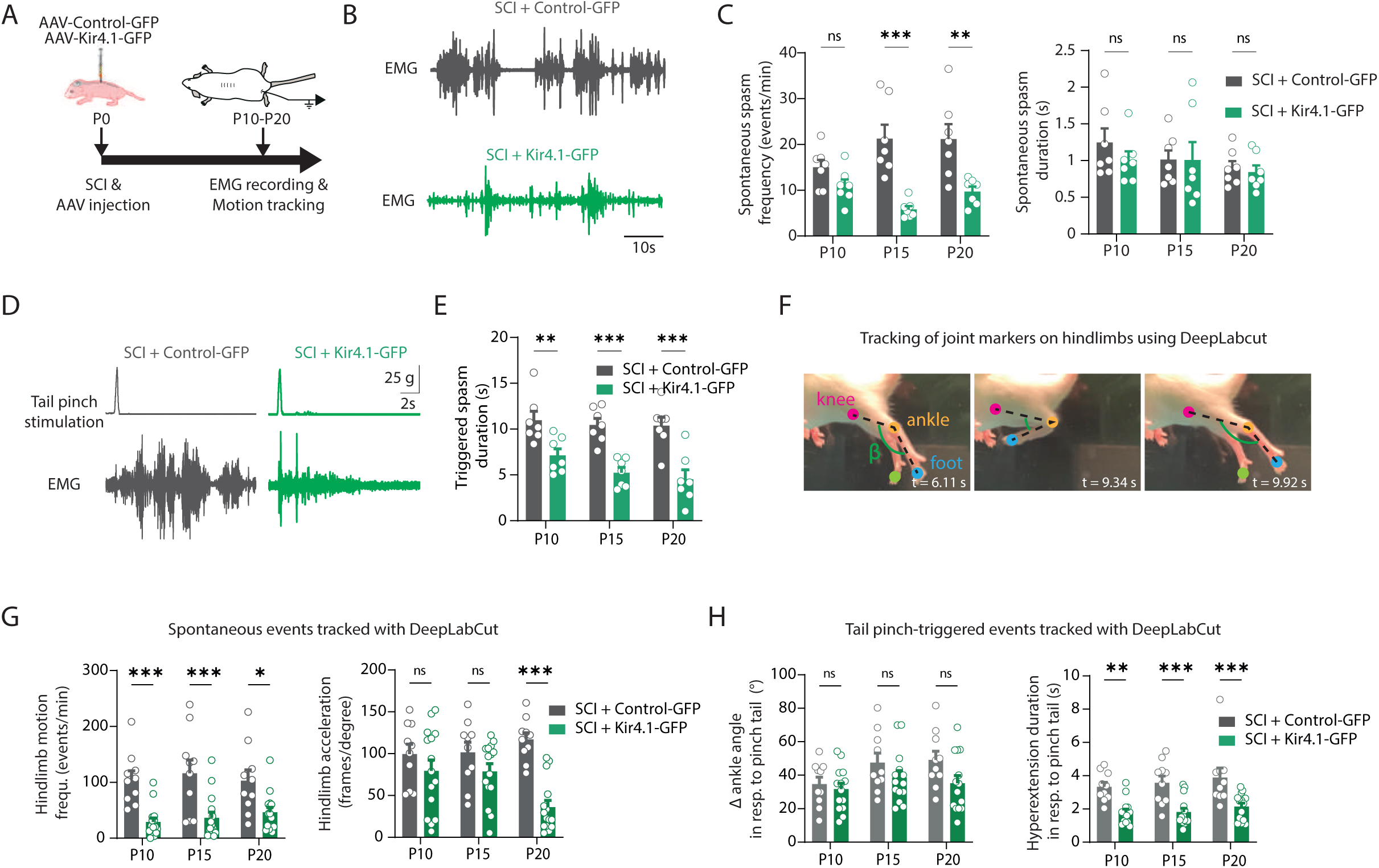
Kir4.1 gain of function in astrocytes reduces spastic-like symptoms in SCI mice. **(A)** Experimental design for AAV targeting astrocyte therapy and *gastrocnemius* EMG recordings. **(B)** Typical *gastrocnemius* spontaneous EMG traces from SCI mice infected with control-AAV (grey) or Kir4.1-AAV (green). **(C)** Quantification of spontaneous muscular event frequency (left) and duration (right) at three different ages in SCI mice injected with control (grey) or Kir4.1-AAV (green). **((D)** Example of *gastrocnemius* EMGs following tail pinch stimulation from SCI mice infected with control-AAV (left, grey) or Kir4.1-AAV (right, green). **(E)** Quantification of tail pinch-evoked muscular event duration at different ages in SCI mice injected with control-AAV (grey) and Kir4.1-AAV (green). For (C-E), SCI + control-AAV: n=7 mice; SCI + Kir4.1-AAV: n=7 mice. **(F)** Example of DeepLabCut tracking of joint markers on hindlimbs. Colored circles show digital markers for analysis. **(G)** Quantification of spontaneous hindlimb motion event frequency (left) and acceleration (right) at different ages in SCI mice injected with control-AAV (grey) or Kir4.1-AAV (green). **(H)** Quantification of ankle angle variation (left) and hindlimb hyperextension duration (right) triggered by tail-pinch stimulation at different ages in SCI mice injected with control-AAV (grey) or Kir4.1-AAV (green). For (G-H), SCI + control-AAV: n=10 mice; SCI + Kir4.1-AAV: n=15 mice. Data: mean ± S.E.M. ns: not significant, *P < 0.05, **P < 0.01, ***P < 0.001, Two-way ANOVA, Sidak’s multiple comparison test. See also Figures S5 and S6.

## Discussion

Our study reveals novel insights into neuro-glial interactions within spinal motor circuits and demonstrate that improving Kir4.1 function in astrocytes reduces spasticity in SCI mice. Employing a multifaceted methodology that includes advanced imaging techniques, electrophysiology, immunohistochemistry, and genetic tools, we explored the effects of enhancing astrocytic K^+^ uptake in alleviating spasticity. We found that thoracic SCI prompts lumbar astrocytes to adopt a reactive phenotype characterized by significant morphological and functional alterations, as well as the upregulation of inflammatory markers. These reactive astrocytes exhibit increased intracellular acidosis mediated by NBCe1, disrupting the function of Kir4.1 channels and leading to compromised K^+^ absorption which heightened MN excitability.

### Morphological and inflammatory changes from lumbar astrocytes following thoracic SCI

Reactive astrocytes undergo morphological, molecular and functional remodelling in response to various CNS injuries in surrounding tissue^23,24,40^. Reactive astrocytes at the glial scar protect tissue and preserve function after SCI by restricting the spread of inflammatory cells^41,42^. However, the occurrence and nature of astrocyte reactivity in regions distal to the injury site post-SCI remain poorly understood. Here, we reveal that thoracic SCI induces significant astrocyte reactivity within the lumbar region, evidenced by hypertrophied and increased complexity of astrocytic processes. We also report an overall rise in GFAP intensity within the ventral horn’s gray matter, without modification of the astrocyte/motoneuron ratio, thus underscoring the pervasive astrocytic presence around α-MNs as observed in amyotrophic lateral sclerosis^43^ or spinal muscular atrophy^44^ mouse models. In our study, we also observed a modest but significant increase in the pro-inflammatory markers (C3d, STAT3) in lumbar astrocytes following SCI. Previous studies focusing on the spinal motor networks, indicated that pro-inflammatory cytokines from microglia, such as IL-1α, IL-6, TNF-α, and C1q, may drive the enhanced astrocytic expression of C3^28,43^ and pSTAT3 signaling^45,46^. Moreover, the persistent activation of astrocytic STAT3 has been shown to depend on increased Ca^2+^ signaling in a mouse model of chronic inflammatory itch^47^. These findings further underscore the significance of exploring the astrocytic Ca^2+^ dynamics after SCI.

### Functional changes from lumbar astrocytes following thoracic SCI

In conjunction with morphological evaluations, we investigated the functional plasticity of lumbar astrocytes post-SCI. Reactive astrocytes demonstrate varied Ca^2+^ dynamics across different pathological models, stages (acute vs. chronic), and anatomical regions^40^. Generally, these cells show dysregulated Ca^2+^ signaling, predominantly characterized by heightened Ca^2+^ activity^48–50^, though reductions in Ca^2+^ signaling have also been reported^51,52^. Our two-photon Ca^2+^ imaging data indicate significant alterations in Ca^2+^ signaling in lumbar astrocytes from SCI mice. Specifically, we observed an increase in both (i) the proportion of spontaneous active astrocytic Ca^2+^ transients and (ii) the glutamate-evoked Ca^2+^ activity in astrocytic soma, processes, and microdomains. Ca^2+^ signaling is crucial as it impacts numerous cellular processes, including the modulation of neurotransmitter release^53^ and K^+^ buffering capacity^54,55^. Moreover, simultaneous Ca^2+^ imaging from both neurons and astrocytes using dual-color genetically encoded calcium indicators (GECIs) revealed that astrocyte-neuron Ca^2+^ signalling is enhanced after SCI. While further studies are needed to elucidate the underlying mechanisms, these findings suggest that the increased and widespread Ca^2+^ signalling in astrocytes may fuel neuronal hyperexcitability and subsequent spasticity following SCI. Supporting this notion, a proconvulsive potential of astrocyte Ca^2+^ signals in promoting the spread of epileptiform activity within the hippocampal networks has been demonstrated^56^.

### pH alterations in lumbar astrocytes post-SCI

Alterations in brain pH are recognized as critical indicators of various pathological states^57^, with astrocytes playing a key role in maintaining an intricate pH balance between the intra and extracellular environments^15^. In our study, we identified two distinct astrocytic populations in control mice that responded to glutamate in divergent ways affecting intracellular pH: the majority (∼70%) exhibited intracellular acidification, likely for maintaining the extracellular pH homeostasis during sustained neuronal activity through activity-dependent HCO3^−^ efflux^33^, whereas the minority (∼30%) showed intracellular alkalinization, possibly facilitated by K^+^-stimulated NBCE1 activity that supports energy metabolism during astrocytic glycolysis^58^. Following SCI, our two-photon imaging data indicated a rise in the proportion of acidifying astrocytes to 92%, with an increase in the amplitude of acidifying pHi responses. These changes could be attributed to a pSTAT3-dependent increase in NBCE1 expression^59^ and a shift in NBCE1 activity towards its outward mode triggered by enhanced astrocytic Ca^2+^ signaling^34,35^. Additionally, chronic hypoxia beneath the injury site, due to compromised capillary blood flow^60^, may further drive NBCe1-induced Ca^2+^ transients in astrocytes, thereby exacerbating intracellular pH acidification^35,61,62^. Additional research is needed to explore the roles of other factors (i.e. NCX, AE, …) in this intensified astrocytic pH acidification^57,63^.

### Acidosis-mediated dysfunction of Kir4.1 leading to increased extracellular K+ post-SCI

Effective extracellular potassium (K^+^) removal is essential for maintaining brain homeostasis and limiting network hyperexcitability, as impairments in K^+^ clearance are associated with various pathological states^38,64–66^. The maintenance of K^+^ homeostasis is predominantly performed by astrocytes, primarily through the activity of the weakly inwardly rectifying Kir4.1 K^+^ channels^21,25^. Given the pH sensitivity of Kir4.1 to acidosis^36,37^, we confirmed that NBCe1-mediated pH acidification impairs Kir4.1 channel function post-SCI, thereby compromising the reactive astrocytes’ capacity to buffer extracellular K^+^. This dysfunction is exacerbated by acidosis but can be reversed by NBCe1 blockade, emphasizing the detrimental impact of acidosis on Kir4.1 activity and resulting K^+^ imbalance post-SCI. We previously demonstrated that SCI displays early and chronic MN hyperexcitability^10,12^, which occurs alongside with the astrocytic Kir4.1 deficits observed in this study. Additionnally, we observed that increase in extracellular K^+^ reduces the delayed rectifying K^+^ current, Kv1.2, potentially amplifying motor output^18^. However, unlike in the hippocampus^67^, the increase in K^+^ did not affect the persistent sodium current (INaP) that is typically amplified post-SCI^11^. This discrepancy may be attributed to the moderate (∼3mM) rise in K^+^ in response to glutamate post-SCI, aligning with the sensory-evoked K^+^ rise observed in striatal and hippocampal networks under pathological conditions^65,66^. Collectively, these findings underscore the importance of correcting K^+^ dysregulation to decreases MN hyperexcitability and thus potentially reduce spasticity post-SCI.

### Kir4.1 gain of function for decreasing spasticity post-SCI

The observation that impaired K^+^ homeostasis leads to MN hyperexcitability post-SCI raise the question of whether increasing astrocytic Kir4.1 function could decrease spasticity. Previous studies have shown that a selective restoration of Kir4.1 function in astrocytes reverts Huntington disease-like symptoms^51,65^, can relieve chronic hyperalgesia in neuropathic pain^68^ and improves cognitive and social performance in a Fragile X syndrome mouse model^66^. Spinal astrocytes might thus play a key role in modulating motor troubles after SCI as: (i) they are present in high proportion in the ventral horn^69^, (ii) they control pathological tremor by tuning excitatory synaptic transmission through the release of purines^19^, and (iii) the Rac1 activity in astrocytes can contribute to hyperreflexia following SCI^70^. Our viral approach specifically targeted Kir4.1 channels, which are highly expressed in astrocytes around large α-MNs^22^.

α-MNs have the capacity to display a sustained firing pattern^71,72^, which is enhanced post-SCI^4,7,12^. By enhancing Kir4.1 function in spinal astrocytes, we observed a significant alleviation of spastic-like symptoms in SCI mice. The reduction in muscle spasm frequency and duration in treated mice highlights the functional importance of Kir4.1 channels in modulating neuronal excitability and controlling spasticity post-SCI.

### Limitations of the Study

Our study provides new biological insights onto neuroglial crosstalk in the spinal motor network and broaden the spectrum of therapeutic targets for restoring compromised neuronal excitability in neurological diseases involving altered K^+^ homeostasis. However, some limitations warrant consideration. Primarily, by focusing on Kir4.1, we may have overlooked the potential contributions of other ion channels or molecular components, notably Na^+^K^+^ ATPase pump, which could play a pivotal role in regulating K^+^ homeostasis^73^. Additionally, in a translational point of view, the juvenile SCI model may not accurately mirror the complexities of adult SCI environment. Moreover, considering the diverse forms of spasticity in humans and the various models used in mice^74^, it will be important in future work to establish whether our findings extend to the other forms and models of spasticity, such as those induced by SCI, stroke or neurodegenerative diseases like Parkinson’s disease or multiple sclerosis. Our results also highlight the significance of astrocyte-selective AAV gene therapy in biomedical research. However, AAV technologies are still in their infancy regarding clinical trials aimed at treating CNS disorders, and optimizing delivery methods to enhance both safety and efficacy^75^ are necessary.

## MATERIALS AND METHODS

### Experimental model

#### Mice and ethical standards

CD1 mice of either sex (P12-P18 for patch-clamp recording, P13-P20 for two-photon Ca^2+^ imaging, P10, P15 and P20 for EMG recordings and DeepLabCut procedure) were housed under a 12h light/dark cycle in a temperature-controlled area with *ad libitum* access to water and food. *Aldh1L1*-eGFP mice (CD1 background) expressing GFP in astrocytes were kindly provided by Nathalie Rouach (Collège de France, CNRS, INSERM, Labex Memolife, Université PSL, Paris, France). Animals from different litters were used for each experiment. Sample sizes (number of spinal cords and/or cells) for each experiment are indicated in figure legends. All animal care and use were conformed to the French regulations and approved by the local ethics committee (Comité d’Ethique en experimentation animale CE n°071 (INT-Marseille, Nb B 13 014 04), APAFIS authorizations: Nb 17485-2018110819197361 and Nb 44572-2023072115506121).

### Experimental procedures

#### Spinal cord injury (SCI)

The spinal cord transsection was performed on P0-P1 neonatal mice, including both wild-type and *Aldh1L1*-eGFP mice. The animals were anesthetized *via* hypothermia and received a subcutaneous injection of buprenorphine (0.025 mg/kg; Vetergesic, CEVA Santé Animale, France). After a midline skin incision, a laminectomy was performed to expose lower thoracic segments of the spinal cord. The dura was opened and the spinal cord was completely transected at the T8-T9 segmental level. The lesion cavity was filled with sterile absorbable collagen contact hemostat (Pangen, Urgo Medical, France). One drop (ca 20-30 µl) of the antibiotic amoxicillin (Citramox, Axience, France) was subcutaneously applied at the incision site. Finally, the wound was closed with 6-0 absorbable suture (Z1032H, Ethicon, NJ) and covered with adhesive suture (Steri-Strip, R1546, 3M Health Care, MN). Animals were kept in a warm and wet chamber for 2 hr in cotton-wool swab impregnated with their mother smell before they returned to the home cage with their mother. Sham animals were submitted to all procedures except the laminectomy and the spinal cord transection.

### Adeno-associated virus (AAV) constructs

AAV2/5-gfaABC1D-cyto-GCaMP6f (52925-AAV5, titer ≥ 1*10^13^ vg/mL), AAV2/9-Syn.NES-jRGECO1a.WPRE.SV40 (100854-AAV9, titer ≥ 1*10^13^ vg/mL), AAV2/5-gfaABC1D-tdTomato (44332-AAV5, titer ≥ 7*10^12^ vg/mL) were sourced from Addgene (https://www.addgene.org/). AAV2/9-GFAP-eGFP-Cre (titer ≥ 2.5*10^10^ vg/mL) AAV2/9-gfaABC1D-eGFP-Kir4.1 (titer ≥ 2.3*10^11^ vg/mL) were kindly provided by the Hu lab^76^.

### Intrathecal AAV delivery

A minimally invasive technique was employed to micro-inject AAV vectors through the intervertebral space. Briefly, in cryoanesthetized pups, a beveled microcapillary (30-60 µm in diameter) preloaded with the AAV particles was lowered to reaches the subarachnoidal space at the L3-L4 segments. A total volume of 2 µL per animal was then progressively injected (1 µL/5s).

### Lumbar slice preparation and artificial cerebro-spinal fluid (aCSF) solution

Mice were anaesthetized with intraperitoneal injection of a mixture of ketamine/xylazine (100mg/kg and 10 mg/kg, respectively). They were then decapitated, eviscerated and the spinal cord removed by laminectomy, and placed in a Sylgard-lined petri dish with ice-cold (1-2°) aCSF containing (in mM): sucrose (252), KCl (3), NaH_2_PO_4_ (1.25), MgSO_4_ (4), CaCl_2_ (0.2), NaHCO_3_ (26), D-glucose (25), pH 7.4. The lumbar spinal cord was mounted on an agar block, embedded in a 4% agarose solution, quickly cooled, and then sliced into 325 µm sections through the L4–L5 lumbar segments using a vibrating microtome (Leica, VT1000S). Slices were immediately transferred into the holding chamber filled with bubbled (95% O_2_ and 5% CO_2_) standard aCSF composed of (in mM): NaCl (120), KCl (3), NaH_2_PO_4_ (1.25), MgSO_4_ (1.3), CaCl_2_ (1.2), NaHCO_3_ (25), D-glucose (20), pH 7.4, 30-32°C. After a 30-60 min resting period, individual slices were transferred to a recording chamber continuously perfused with standard aCSF heated to 32-34°C.

### *Ex vivo* electrophysiological recordings

Aldh1L1-eGFP positive astrocytes were visualized in the ventrolateral region of lamina IX in L4-L5 horizontal slices using an Nikon FN1 microscope with appropriate filters. Whole-cell patch-clamp recordings of GFP(+) astrocytes and surrounding large motoneurons (MNs) (soma area >800 µm^2^)^72^ were performed using a Multiclamp 700B amplifier (Molecular Devices) with electrodes pulled from borosilicate glass capillaries (1.5 mm OD, 1.12 mm ID; World Precision Instruments) on a Sutter P-97 puller (Sutter Instruments Company). For MN recordings, electrodes (2-4 MW) were filled with an intracellular solution containing (in mM): K^+^-gluconate (140), NaCl (5), MgCl_2_ (2), HEPES (10), EGTA (0.5), ATP (2), GTP (0.4), pH 7.3. For GFP+ astrocyte recordings, electrodes (6-8 MW) were filled with an intracellular solution containing (in mM): K^+^-gluconate (105), NaCl (10), KCl (20), MgCl_2_ (0.15), HEPES (10), EGTA (0.5), ATP (4), GTP (0.3), pH 7.3. In some experiments, the intracellular pH was lowered to 6.9 by the addition of HCl. Patch clamp recordings were made using a Multiclamp 700B amplifier driven by PClamp 10 software (Molecular Devices). Recordings were digitized on-line and filtered at 10 kHz (Digidata 1550B, Molecular Devices). Pipette and neuronal capacitive currents were canceled and, after breakthrough, the access resistance was compensated. All experiments were designed to gather data within a stable period (i.e., at least 1-2 min after establishing whole-cell access).

### *Ex vivo* two-photon Ca^2+^ imaging

Two-photon fluorescence measurements were obtained with a dual-scanhead two-photon microscope (FemtoS-Dual, Femtonics Ltd, Budapest, Hungary) and made using an Olympus XLUMPlanFLN 20X, 1.00 NA objective (Olympus America, Melville, NY) or a Nikon LWD 16X/0.80W objective (Nikon Instruments Inc. Melville, NY). For two-photon Ca^2+^ imaging, excitation of GCaMP6f and jRGECO1a was simultaneously evoked with a femtosecond pulsed laser (Chameleon Ultra II; Coherent, Santa Clara, CA) tuned to 960 nm. Recordings were performed at ∼32−34 °C in standard aCSF saturated with 95% O_2_/5% CO_2_ (pH 7.4). The microscope system was controlled by MESc acquisition software (https://femtonics.eu/femtosmart-software/, Femtonics Ltd, Budapest, Hungary). A single acquisition plane was selected and full-frame imaging was started in a resonant scanning mode at 30.5202 Hz. Scan parameters were [pixels/line × lines/frame (frame rate in Hz)]: [512 × 519 (30.5202)]. Scanning area was 468 µm * 474 µm. This microscope was equipped with two detection channels for fluorescence imaging.

### *Ex vivo* two-photon imaging of intracellular pH (pH_i_) in astrocytes

Lumbar slices were incubated in standard aCSF containing BCECF (5 µM) and Pluronic® F-127 (0.02%) for ≃ 45 min followed by a through washing out to allow de-esterification of the dye. Two-photon fluorescence measurements were obtained with a dual-scanhead two-photon microscope (FemtoS-Dual, Femtonics Ltd, Budapest, Hungary) and made using an Olympus XLUMPlanFLN 20X, 1.00 NA objective (Olympus America, Melville, NY) or a Nikon LWD 16X/0.80W objective (Nikon Instruments Inc. Melville, NY). For two-photon BCECF signal imaging, excitation was simultaneously evoked with a femtosecond pulsed laser (Chameleon Ultra II; Coherent, Santa Clara, CA) tuned to 820 nm. Recordings were performed at ∼32−34 °C in aCSF saturated with 95% O_2_/5% CO_2_ (pH 7.4). Astroglial loading of BCECF was confirmed by analysing the characteristic morphology of the BCECF-stained cells featuring astrocytic phenotype from mice injected at birth with AAV AAV2/5-gfaABC1D-tdTomato. Changes in pH*_i_* are expressed as changes in BCECF fluorescence at the maximum of the fluorescent signal over the baseline (Δ*F*/*F*_0_). The microscope system was controlled by MESc acquisition software (https://femtonics.eu/femtosmart-software/, Femtonics Ltd, Budapest, Hungary). A single acquisition plane was selected and full-frame imaging was started in a resonant scanning mode at 30.5202 Hz. Scan parameters were [pixels/line × lines/frame (frame rate in Hz)]: [512 × 519 (30.5202)]. Scanning area was 468 µm * 474 µm. This microscope was equipped with two detection channels for fluorescence imaging.

### Puff-application of extracellular K^+^

To measure the astrocytes’ potassium (K+) uptake capacity, a high extracellular potassium concentration ([K^+^]_o_, 12mM) was added to standard aCSF and delivered via a pipette similar to the recording pipette, controlled by a Picospritzer (Picrospritzer II, General Valve Corporation) set at 10 psi. The electrode was positioned in the ventro-lateral part (lamina IX) of a lumbar slice, within ∼60 μm of a GFP+ astrocyte soma.

### Measurement of extracellular K^+^ concentration

K^+^-sensitive microelectrodes were made from thin-walled borosilicate theta tubes (GC150T-10, Harvard Apparatus, MA) following the procedure described in^77^. Briefly, the theta tubes were washed in 70% ethanol for 1 hour, then for 1 hour in ultra-pure water, and dried in the oven overnight at 60°C. The double-channel pipettes were pulled to obtain a tip diameter of 3-5 µm. One channel was filled with 150 mM NaCl and the second one with 100 mM KCl. The inside of the tip of the KCl-containing channel was silanized using Silanization Solution I (85126, Merck-Sigma Aldrich) and then filled with Potassium Ionophore I Cocktail A (99311, Merck-Sigma Aldrich). The electrode tip was immersed in a 100 mM KCl solution for 3 hours or longer for stabilization. Before each recording session, the electrode was calibrated using ACSF solutions enriched with gradually decreasing KCl concentrations: 20 mM, 10 mM, 6 mM, 3 mM, 1.25 mM, 0.6 mM, 0.3 mM. The K^+^ sensor was accepted for the experiment if it generated a stable potential at each K^+^ concentration and displayed a >540 mV increase at 30 mM K^+^ drop.

The extracellular potassium concentration ([K^+^]_o_) was measured in acute lumbar slices from control and spinal cord injury (SCI) mice. Experiments were conducted in standard aCSF, saturated with 95% O2/5% CO2 (pH 7.4) and heated to 32-34°C. L-glutamate (1 mM, 5 s, at 10 psi) was puffed using a glass pipette connected to a picospritzer (Picrospritzer II, General Valve Corporation). Both the puff electrode and the K^+^-sensitive microelectrode were aligned and positioned in the ventro-lateral part (lamina IX) of the lumbar slice, within ∼50 μm of each other in the z axis. Basal [K^+^]_o_ was measured after proper positioning and stabilization of the K^+^-sensitive microelectrode in the lumbar slice.

Data acquisition was performed in current-clamp mode using a Multiclamp 700B amplifier (Molecular Devices), sampled at 10 kHz, filtered at 4 kHz, and digitized with a Digidata 1550B (Molecular Devices). The peak amplitude, area under the curve (AUC) of [K^+^]_o_ transients, the rise slope from glutamate puff onset to peak amplitude, and decay time from peak amplitude to baseline recovery were measured.

### Pharmacological compounds and dyes

All solutions were oxygenated with 95% O_2_/5% CO_2_. Barium chloride dihydrate (Ba^2+^, 100µM; 217565), Pluronic® F-127 (0.02%; P2443), DMSO (0.1%; D4540), L-glutamic acid (L-glutamate, 1 mM; G1251) and Sodium Propionate (5mM; P1880) were obtained from Merck Sigma-Aldrich. Tetrodotoxin (TTX, 0.5-1µM; 1078) from Tocris Bioscience. S0859 (50-100 µM, M5069) from Abmole Europe Branch (Forlab). 2’,7’- Bis-(2-Carboxyethyl)-5-(and-6)-Carboxyfluorescein, Acetomxymethyl Ester (BCECF-AM, 5 µM) from Invitrogen. All drugs were dissolved in water and added to the standard aCSF except BCECF-AM incubated in Pluronic® F-127 (0.02%) and DMSO (0.2%).

### Immunohistochemistry

Spinal cords of 14-16 days-old mice were.dissected out and fixed for 5-6 h in 4% paraformaldehyde (PFA), then rinsed in phosphate buffered saline (PBS) and cryoprotected overnight in 20% sucrose at 4°C. Spinal cords were frozen in OCT medium (Tissue Tek) and 30 μm cryosections were collected from the L4-L5 segments. After having been washed in PBS 3×5 min, the slides were incubated for 1 h in a blocking solution (BSA 1%, Normal Donkey Serum 3% in PBS) with 0.2% triton X-100 and for 24 h at 4 °C in a humidified chamber with the following primary antibodies diluted in the blocking solution with triton x-100: rabbit anti-Kir4.1 (1/1000; APC-035, Alomone, Israel), mouse anti-NeuN (1/1000; MAB377, Merck Millipore, MA), rabbit anti-GFAP (1/1000; Z0334, Dako, Agilent, CA) and/or goat anti-C3d (1/40; AF2655, R&D Systems, MN). For STAT-3 staining, slices were pretreated with a drop of MetOH 100% (34860-1L-R; Merck Sigma-Aldrich, MA) for 10 minutes at -20°C. After 3 washes of 5 min in PBS, they were incubated for 1h in a blocking solution (Normal Donkey Serum 5% in PBS 0.1M) with 0.3% triton X-100. The primary antibody anti-STAT3 (1/300; 8768, Cell Signaling Technology, MA) was incubated with Cell signal blocking solution (8112, Cell Signaling Technology, MA) for 48 h at 4°C. Slides were then washed 3×5 min in PBS and incubated for 2 h with the appropriate secondary antibodies diluted in the blocking solution: Alexa Fluor® Plus 555-donkey anti-rabbit (1/400; A32794, Invitrogen, MA), Cy5-donkey anti-mouse (1/400; 715-175-151, Jackson ImmunoResearch, PA) and/or Alexa Fluor® Plus 555-donkey anti-goat (1/400; A32816, Invitrogen, MA). After 3 washes of 5 min in PBS, slides were incubated with Hoechst 33342 (2 µM in PBS; 62249, Thermo Fisher Scientific, MA) for 5 min and washed again 3x5 min before being mounted with a homemade gelatinous aqueous medium. Images were acquired using a confocal microscope (LSM700, Zeiss) equipped with either a 20x air objective, a 40x oil objective or a 63x oil objective and processed with the Zen software (Zeiss).

### Protein analysis by capillary Western Blot

The lumbar parts of the spinal cords were dissected in aCSF at 4°C and conserved at -80°C until protein extraction. Tissues were homogenized in ice-cold lysis buffer (250 mM sucrose, 3.9 mM Tris pH 7.5, 10 mM iodoacetamide) supplemented with protease inhibitors (cOmplete™, Mini, EDTA-free Protease Inhibitor Cocktail, 11836170001, Roche Diagnostics, Germany). Unsolubilized material was pelleted by centrifugation at 7,000 x g for 5 min at 4°C and discarded. The supernatants were subjected to an additional centrifugation step at 19,000 x g for 70 min at 4°C. The resulting pellets corresponding to the membrane-enriched fraction were resuspended in ice-cold lysis buffer (PBS 1X, IGEPAL® CA-630 1%, SDS 0.1%) supplemented with protease inhibitors (cOmplete™, Mini, EDTA-free Protease Inhibitor Cocktail, 11836170001, Roche Diagnostics, Germany). Protein concentrations were determined using the Pierce™ BCA Protein Assay Kit (23227, Thermo Fisher Scientific, MA). Kir4.1 and NBCe1 expressions were then analyzed using the 12-230 kDa and the 66-440 kDa separation modules respectively (SM-W004 and SM-W008, ProteinSimple, Bio-Techne, MN) on an automated capillary western blotting system (‘Jess’, ProteinSimple, Bio-Techne, MN) according to the manufacturer’s protocol with small modifications. Samples were not implemented with DTT and they were not submitted to heat denaturation in order to keep proteins in a more ‘native’ form. Total protein concentrations of 0.1 mg/mL and 0.5 mg/mL were used for Kir4.1 and NBCe1 respectively. Samples were probed with a rabbit polyclonal Kir4.1 antibody (1:60; APC-035, Alomone, Israel) or a rabbit polyclonal NBCe1 antibody (1:90; ANT-075, Alomone, Israel) and revealed with the appropriate detection module (DM-001, ProteinSimple, Bio-Techne, MN). Loaded samples were normalized to their own total protein content using the ‘total protein detection module’ (DM-TP01, ProteinSimple, Bio-Techne, MN).

### Assessement of electromyography (EMG) recordings

Mice were tested at 10, 15, and 20 days post-SCI, when signs of spastic motor behaviors, such as excessive involuntary twitches/movements and exaggerated reflexes, were evident. For electrophysiological assessment of spasticity, a stainless steel needle electrode was transcutaneously inserted into the lateral gastrocnemius muscle (ankle extensor). Two reference electrodes were placed subcutaneously on the back and in contact with the patellar tendon. Because mice were spinalized, they did not feel any pain during this procedure. Following a 30-minute acclimation period to the recording environment, mice were immobilized with tape. Spontaneous motor responses were recorded for 15 minutes. Motor responses evoked by tail pinch (50.5 ± 5.6 g, 0.5 ± 0.2 s) were recorded until baseline activity returned. Each mouse experienced five consecutive sensory stimuli, each separated by a resting period of at least 30 seconds. The proximal ends of the recording wires were connected to a A-M Systems Amplifier (Model 1700, Everett, WA). EMG signals were amplified (1000x) and bandpass filtered (100 Hz to 5 kHz), then sampled at 13.5 kHz (Digidata 1440A, Molecular Devices). The applied pressure (measured in grams) to the distal third of the tail was recorded by a miniature pressure sensor (#FS1901-000-0500-G, Farnell) placed between the thumb and the tail, and monitored in real-time.

### Assessment of DeepLabCut video acquisition

Animals were tested at P10, P15, and P20, when signs of a spastic motor phenotype, such as excessive involuntary hindlimb twitches and exaggerated reflexes, were visible.

#### Assessment of mice hindlimb spontaneous activity

Animals were removed from their home cages, weighted, and individually subjected to a single 20-second video recording in a standardized homemade setup. Video acquisition was performed at 240 frames per second (FPS) to allow complete visualization of hindlimb spasms from a lateral view, at a distance of 10 cm from the mouse.

#### Assessment of mice hindlimb induced activity

For the behavioral assessment of induced activity, the same procedure was applied as previously described. Induced activity was initiated by gently pinching the tail (without causing discomfort or signs of physical pain) with the thumb. Hindlimb motor responses following tail pinching were recorded by a miniature pressure sensor placed between the thumb and the tail and monitored online. Each animal underwent three consecutive pinching stimulations, separated by 10 seconds, through a single video recording

### Experimental design and statistical analysis

No statistical method was used to predetermine sample size. Group measurements are expressed as means ± S.E.M. The Mann-Whitney test was used for comparisons between two groups, while the Fisher’s test was employed to compare percentages. For comparisons between two conditions, the Wilcoxon matched pairs test was utilized. One-way or two-way ANOVA test was applied for multiple comparisons. Normality of the data sets was evaluated for all statistical analyses, and the significance level was set at p < 0.05. Statistical analyses were conducted using GraphPad Prism 9 software. Details of the analyses are provided in each figure legend.

### Data analysis

#### Assessment of astrocyte reactivity

i. ***Morphometric analyses.*** Confocal images at 63X magnification, immunostained with GFAP antibody in Aldh1L1-eGFP mice, were serially stacked and maximally projected. The maximal projection image of the GFAP signal in the ventral horn of the spinal cord was used for analysis. The maximal projection image of the GFAP signal in the ventral horn of the spinal cord was used for analysis. The Sholl analysis plugin in ImageJ automatically drew serial concentric circles at 5 μm intervals from the center of the nucleus (Hoechst signal) to the end of the longest process in each individual astrocyte. This plugin provided key morphometric parameters such as the number of intercepts of GFAP processes with each circle, the ramification index, and the ending radius (longest process). Four different fields from the left and right parts of the ventral spinal cord were analyzed in at least five different slices from the same mouse, evaluating all astrocytes within the field. For illustrative purposes, a few representative astrocytes were fully reconstructed in 3D using Chimera software.
ii. **Evaluation of C3d positive astrocytes:** Immunofluorescence staining was analyzed from stacked confocal images (15 steps; Z-step, 1 μm, maximum intensity stack) acquired with a 40× objective on five slices per mouse, with four different fields on each slice. All Aldh1L1-eGFP positive astrocytes in the field, including those co-expressing C3d markers (Aldh1L1+ C3d+), were evaluated.
iii. **Quantification of astrocytic STAT3 nuclear fraction.** To evaluate the nuclear fraction of the STAT3 signal, we measured the STAT3 signal intensity within the Hoechst (nuclei) signal. Immunofluorescence staining was analyzed from stacked confocal images (15 steps; Z-step, 1 μm, maximum intensity stack) acquired with a 40× objective on five slices per mouse, with four different fields on each slice. All Aldh1L1-eGFP positive astrocytes in the field with a clear Hoechst signal were included. Nuclear masks were manually segmented using the Hoechst signal, applied to the STAT3 channel, and the mean fluorescence intensity was measured (see ^30^).

### Analysis of *ex vivo* electrophysiological (Patch-clamp) recordings

Electrophysiological data were analyzed offline using Clampfit 10.7 software (Molecular Devices). To ensure optimum quality of intracellular recordings, several criteria were established: only cells exhibiting a stable resting membrane potential (RMP), access resistance with less than 20% variation, and an action potential amplitude larger than 40 mV under standard aCSF were considered.

#### Passive Membrane Properties

Passive membrane properties of astrocytes and motoneurons (MNs) were measured by determining the largest voltage deflections induced by small current pulses from the holding potential, avoiding activation of voltage-sensitive currents. Input resistance was determined by the slope of linear fits to voltage responses evoked by small positive and negative current injections.

#### Recording Kir4.1-Currents in GFP (+) Astrocytes

Voltages were repeatedly stepped for 2.5s from -140 to 0 mV in 10 mV increments, with astrocytes initially clamped at their RMP. Approximately 5 minutes after the start of the recording, barium chloride (Ba^2+^, 100 µM) was bath-applied to the slice. The current amplitude was measured as the difference between the baseline level before the initial voltage step and the mean amplitude over a 10 ms window starting 10 ms after the onset of the hyperpolarizing voltage step. The Ba^2+^-sensitive current was obtained by subtracting the current after Ba^2+^ application from the current before application.

#### MNs excitability

For MNs, firing properties were measured from depolarizing current pulses of varying amplitudes. The rheobase was defined as the minimum step current intensity required to induce an action potential from the membrane potential held at RMP. The instantaneous discharge frequency was determined as the inverse of the interspike interval. The firing gain refers to the ratio between the measured instantaneous firing frequency (Hz) and the injected current (pA).

### Analysis of two-photon Ca^2+^ and BCECF pH_i_ changes imaging

The fluorescent time series depicting Ca^2+^ and BCECF changes in ex vivo neurons and/or astrocytes were analyzed using MESc data acquisition software (Femtonics Ltd, Budapest, Hungary) and the MES curve analyzer tool. Regions of interest (ROIs) for neurons and astrocytic soma were selected manually, while ROIs for astrocytic processes and microdomains were selected automatically using a custom script in ImageJ (Fiji). A signal was declared as a Ca^2+^ transient if it exceeded the baseline by more than twice the baseline noise (standard deviation, SD). We defined active astrocytes as astrocytes displaying at least one Ca^2+^ transient during the recording session.

In Ca^2+^ imaging experiments, ROIs included : (i) active soma for neurons (jRGECO1a), (ii) active soma, processes and microdomains for astrocytes (GCaMP6f). In BCECF experiments, ROIs corresopnded to astrocytic soma double-labeled with Td-tomato (AAV2/5-gfaABC1D-TdTomato).

For the two-photon Ca^2+^ imaging analysis, raw fluorescence traces were extracted to Excel and analyzed using a custom Matlab script. These traces were converted to show fluorescence changes according to Equation (1):

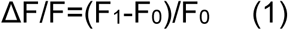

where *F* is the fluorescence at any given time, and F_0_ is the mean fluorescence value for the 5 to 10 s range, preceding the bath perfusion of L-Glutamate (1 mM). The signal-to-noise ratio (SNR) was calculated as:

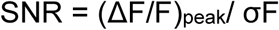

where σF is the SD of the baseline period. Events were considered as Ca^2+^ response if they exceeded twice the SD of the baseline period. The peak amplitude, duration and frequency of both spontaneous and glutamate-evoked Ca^2+^ transients were analysed by using a Matlab script.

For the two-photon pH_i_ imaging analysis, raw fluorescence traces were extracted to Excel. The proportion of Tdtomato (+) astrocytes showing increased or decreased fluorescence (ΔF/F) and the peak amplitude of acidic pHi reponses to L-glutamate were analyzed from both control and SCI mice.

### Analysis of *in vivo* electromyography (EMG) recordings

Custom-built amplifiers enabled simultaneous online rectification and integration (100ms time constant) of the raw AC-mode signals. Data analysis was performed offline with the automatic event detection plugin in Clampfit 11 software. Spontaneous muscle contractions were identified when the signal envelope exceeded a predetermined threshold, and the contraction was considered to end when the signal fell below this threshold. This threshold was adjusted for each signal, typically set at 1.3 times the background noise level. For an event to be classified as a valid muscle spasm, it needed to remain above the threshold for at least 250 ms. We quantified both the average and cumulative distribution of the duration and frequency of these muscle events over a 15 min intervals. For responses triggered by tail stimulation, spasm duration was measured from the stimulus artefact until the recovery of a stable baseline. The experimenter was blinded during both the procedure and the data analysis of the electrophysiological experiments.

### Analysis of mice spastic motor behaviors with DeepLabCut

#### Analysis of kinematics hindlimbs spontaneous activity

For hindlimb spams analysis, videos captured during experiments were analyzed using the markerless pose estimation software DeepLabCut 2.2 (DLC)^39^ to extract tracks based on digital markers placed on the mice (Figure 7). For tracking DeepLabCut was installed on a PC equipped with a NVIDIA Ge-Force RTX 3060 12Go graphics card. DeepLabCut was used to identify four reference points among the mouse body (hip, knee, ankle and foot) on a side view (Figure 7) for kinematic analysis. The ResNet-50 based neural network (imgaug augmentation, 500,000 iterations) was used to identify points of interest on the mice. The training was performed on a set of 800 frames randomly selected by DeepLabCut from 40 videos representatives of experimentation conditions (same background, luminosity, distance from the mouse and the camera).

Note that two separate training has been performed between the first post-natal and second post-natal week face to the very fast morpho-physical changes occurring in mice at this period. Network was trained until it reaches a stable error rate plateau (> 400,000 iterations, 0.0005 error per frame). Next network was evaluated (test error = 2.89 pixels 0,19 mm), and videos analysis was consecutively performed. Raw data provide by DeepLabCut was extracted and proceeded to collect key parameters such as the number of events, acceleration of the spasms, maximum angle of the ankle and the mean ankle angle using custom scripts in Python (see below).

i. *Acceleration*. Sudden increase in the speed of the hindlimb was quantify by estimating the instantaneous acceleration derived from the instantaneous speed estimates:

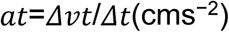

(ii) *Number of events.* Hindlimbs motion event was considered as ankle angle variation of more than 10° degree from the initial position through the entire video recording

### Analysis of hindlimb evoked activity

*(i) Measure of the ankle angle variation following stimulation.* Angle of the ankle was measured using the angle tool of Fiji (ImageJ) before stimulation and immediately following the stimulation. Each measure is performed for the 3 consecutives stimulation for each animal.
*(ii) Measure of spasms duration.* Spasms duration was measured from the end of the stimulation until the recovery of a stable hindlimb motion activity (e.g no hindlimb twitches) using the free and open source cross-platform multimedia player and framework, VLC.
*(iii) Measure of tail pinch force*. Stimulation intensity was recorded by a miniature pressure sensor (#FS1901-000-0500-G, Farnell) placed between the thumb and the tail, and monitored in real-time. The applied pressure measurements were compared between the groups (SCI + control-GFP *vs* SCI + Kir4.1-GFP) and were similar.

### Data Source File

https://github.com/remibos/Data_Source_File_Barbay_SCI_Kir4.1

### Code/software

MATLAB scripts :

https://github.com/remibos/Analysis_2P.git

Python script :

https://github.com/remibos/Analysis_Spasms_DLC

## Data and code availability

- All data reported in this paper will be shared by the lead contact upon request
- All original codes have been deposited at https://github.com/remibos/Analysis_2P.git or https://github.com/remibos/Analysis_Spasms_DLC and are publicly available as of the date of publication.
- Any additional information required to reanalyse the data reported in this paper is available from the lead contact upon request

### Acknowledgments

We thank A. Duhoux for animal care, I. Vanzetta, and A. Lombardini for their technical advices in imaging sessions. J. Verneuil for kindly providing his original scripts. We also gratefully acknowledge E. Gascon, F. Jaouen, C. Lepolard and A. Borges-Correia from Neuro-Vir platform of the Neuro-Bio-Tools facility (Institut de Neurosciences de la Timone, UMR 7289, Marseille, France) for their advices, support and assistance in the design and production of vectors used in this work. We also thank L. Ben Haïm for kindly providing the protocol for STAT3 staining/analysis, and G. Rougon for her critical insights on the manuscript.

## Author contributions

Research design, T.B., E.P. and R.B..; Methodology, T.B., E.P., J.J.R.F., A.I. and R.B.; Investigation and data analysis, T.B., E.P., J.J.R.F. and R.B.; Figures design, T.B. and R.B.; Consulting/Editing, N.R., F.B.; Writing, T.B. and R.B.; Funding Acquisition, R.B.; Supervision, R.B.

## Declaration of interests

The authors declare no competing interests.

## Inclusion and diversity

We support inclusive, diverse, and equitable conduct of research.

## Supplemental information

Document S1. Figures S1–S6

**Figure S1:** Increased GFAP staining is associated with increased inflammatory markers the lumbar segments after thoracic SCI. Related to Figure 1.

**Figure S2:** Spontaneous and glutamate-evoked Ca^2+^ transients in astrocytic soma and microdomains after SCI. Related to Figures 2 and 3.

**Figure S3:** Effect of propionic acid on astrocytic intracellular pH monitored with with the pH-sensitive probe BCECF. Related to Figure 4.

**Figure S4:** Blockade of Kir4.1 channels affects astrocytic K^+^ uptake. Related to Figure 5.

**Figure S5:** Efficiency, specificity and diffusion of the viral strategy for boosting astrocytic Kir4.1. Related to Figure 7.

**Figure S6:** Complementary metrics for assessing spasticity using EMG recordings and DLC tracking. Related to Figure 7.

## Funding sources

This work was mainly funded by the CNRS, the “Fonds d’Investissement de l’INT jeunes chercheuses, jeunes chercheurs” (FI_INT_JCJC_2021) (to R.B.), by the French Ministry of Higher Education, Research and Innovation (PhD fellowship 2021-2024 to T.B.) and by the Network Glia e.V. (Travel grant to TB).

## Notes

### Competing Interest Statement

The authors have declared no competing interest.

