## Supplemental Figures for "Functional contribution of astrocytic Kir4.1 channels to spasticity after spinal cord injury"

Figure S1

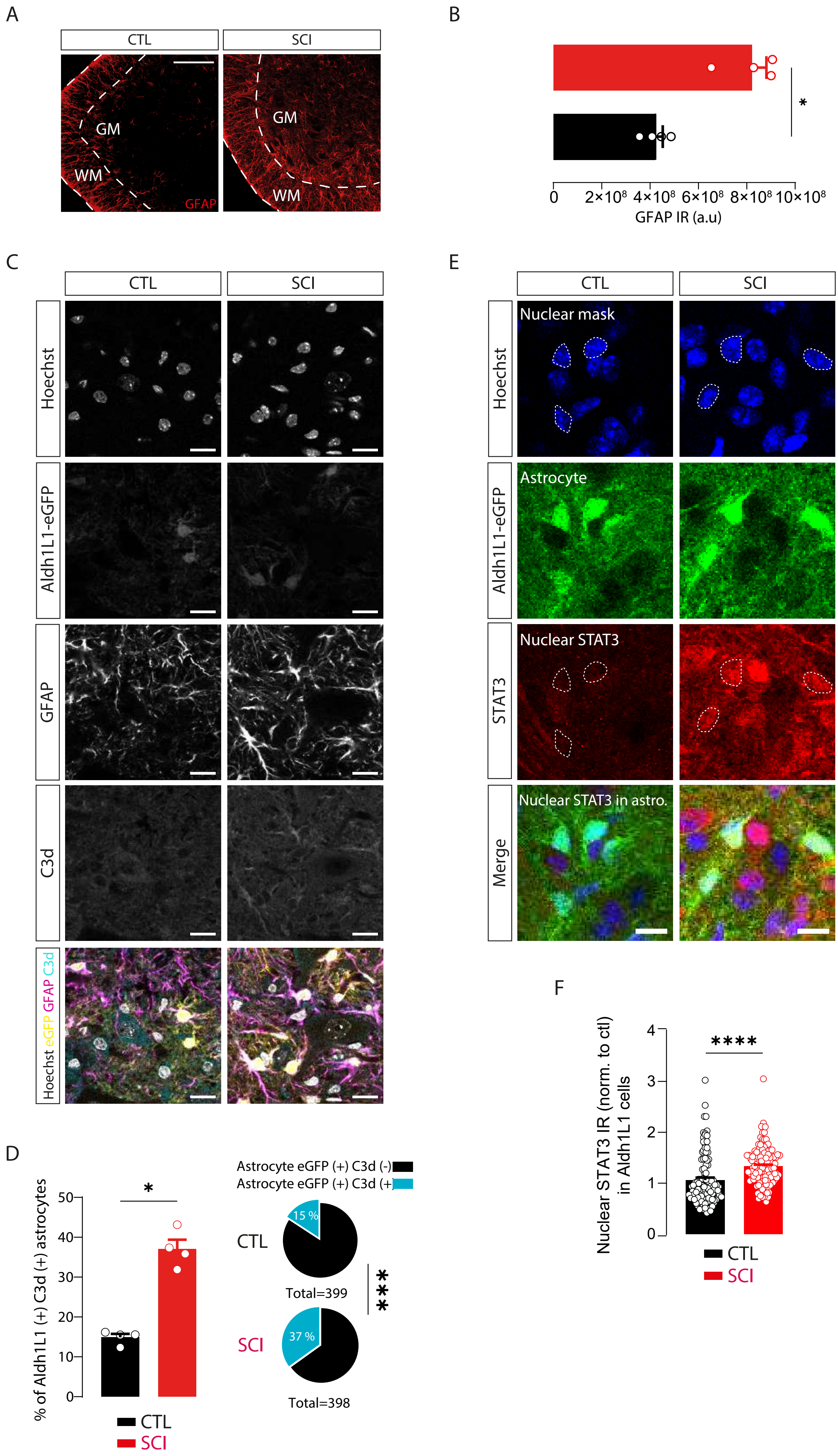

A
CTL
SCI
GM
WM
GFAP
B
*
0
2×10⁸
4×10⁸
6×10⁸
8×10⁸
10×10⁸
GFAP IR (a.u)
C
CTL
SCI
Hoechst
Aldh1L1-eGFP
GFAP
C3d
Hoechst eGFP GFAP C3d
D
*
% of Aldh1L1 (+) C3d (+) astrocytes
50
40
30
20
10
0
CTL
SCI
Astrocyte eGFP (+) C3d (-)
Astrocyte eGFP (+) C3d (+)
CTL
15 %
Total=399
***
SCI
37 %
Total=398
E
CTL
SCI
Hoechst
Nuclear mask
Aldh1L1-eGFP
Astrocyte
STAT3
Nuclear STAT3
Merge
Nuclear STAT3 in astro.
F
****
Nuclear STAT3 IR (norm. to ctl) in Aldh1L1 cells
4
3
2
1
0
CTL
SCI

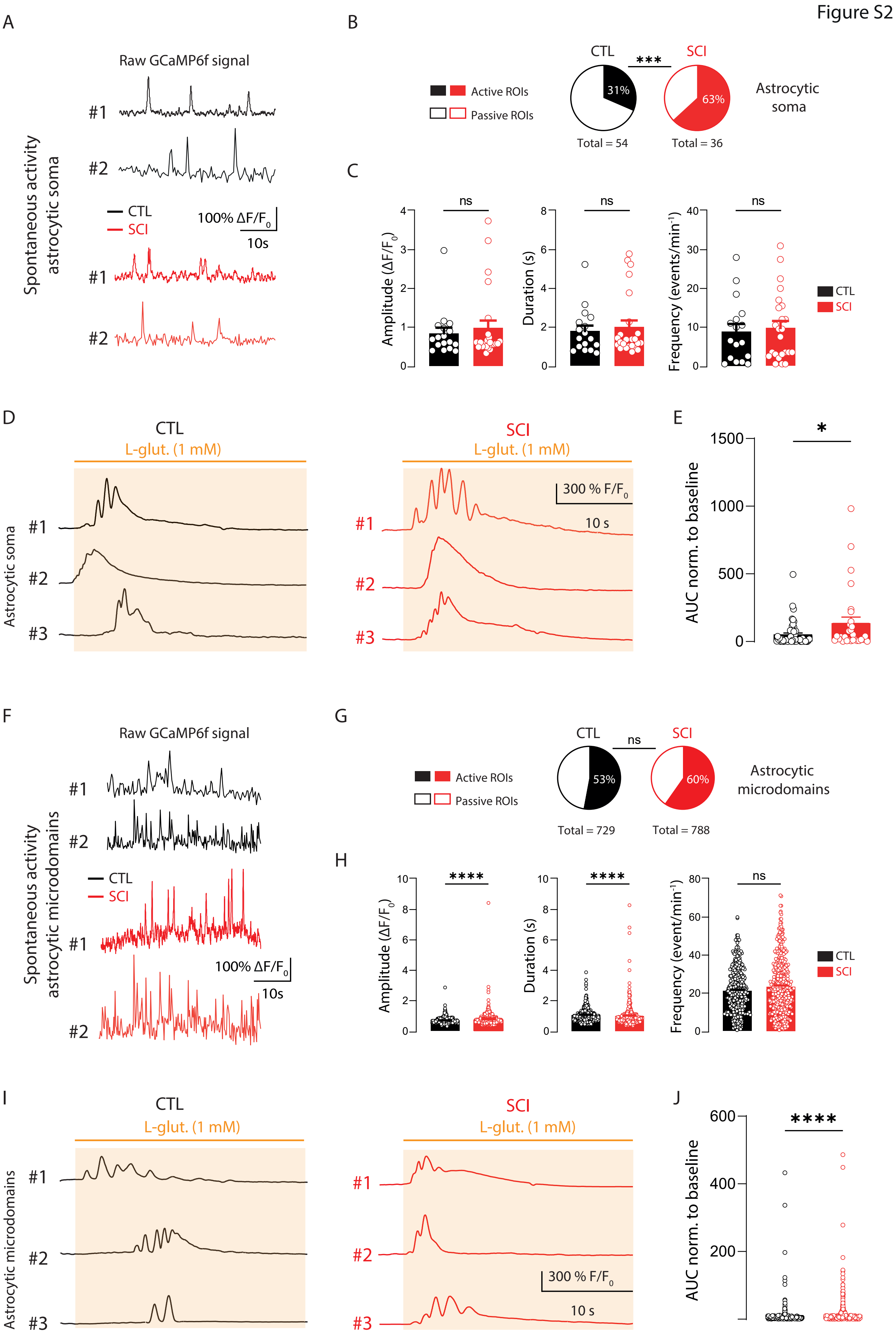

Figure S2
A
Raw GCaMP6f signal
Spontaneous activity astrocytic soma
#1
#2
CTL
SCI
100% ΔF/F0
10s
B
CTL
***
SCI
Active ROIs
Passive ROIs
31%
63%
Total = 54
Total = 36
Astrocytic soma
C
Amplitude (ΔF/F0)
Duration (s)
Frequency (events/min-1)
ns
D
CTL
SCI
L-glut. (1 mM)
Astrocytic soma
#3
300 % F/F0
10 s
E
AUC norm. to baseline
*
F
Spontaneous activity astrocytic microdomains
G
ns
53%
60%
Total = 729
Total = 788
Astrocytic microdomains
H
****
Frequency (event/min-1)
I
Astrocytic microdomains
J
****

Figure S3

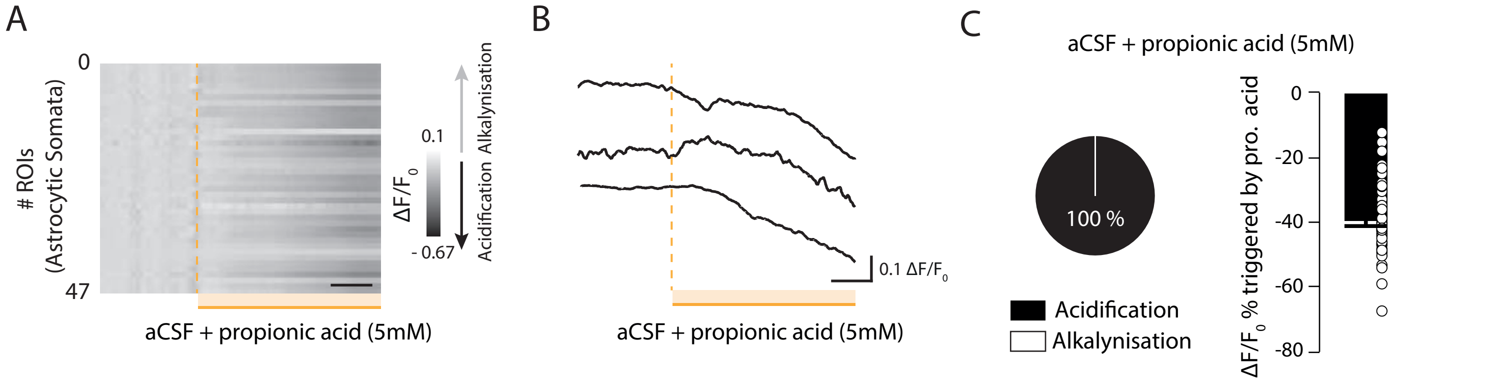
A
0
47
### ROIs
(Astrocytic Somata)
0.1
- 0.67
ΔF/F0
Acidification Alkalynisation
aCSF + propionic acid (5mM)
B
0.1 ΔF/F0
aCSF + propionic acid (5mM)
C
aCSF + propionic acid (5mM)
100 %
Acidification
Alkalynisation
ΔF/F0 % triggered by pro. acid
0
-20
-40
-60
-80

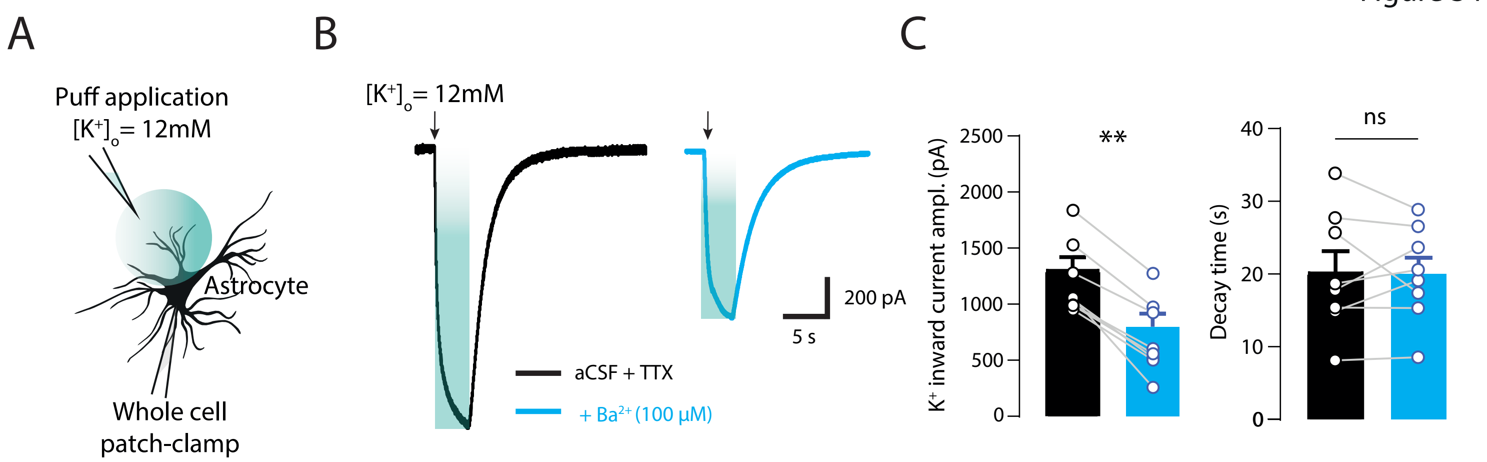
A
Puff application
[K+]o = 12mM
Astrocyte
Whole cell
patch-clamp
B
[K+]o = 12mM
200 pA
5 s
aCSF + TTX
+ Ba2+ (100 µM)
C
K+ inward current ampl. (pA)
2500
2000
1500
1000
500
0
**
Decay time (s)
40
30
20
10
0
ns

Figure S5

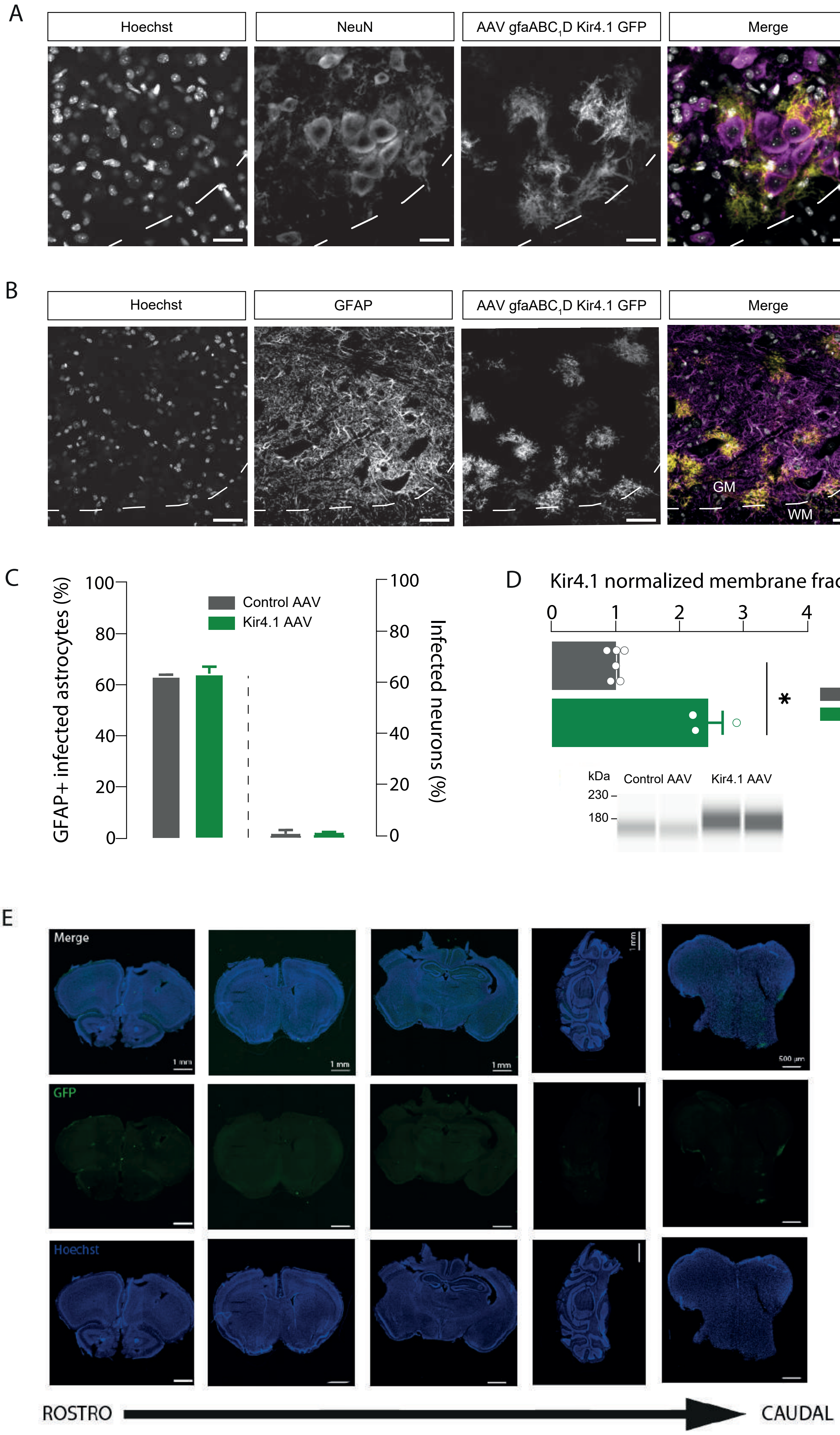
A
Hoechst
NeuN
AAV gfaABC1D Kir4.1 GFP
Merge
B
Hoechst
GFAP
AAV gfaABC1D Kir4.1 GFP
Merge
GM
WM
C
GFAP+ infected astrocytes (%)
100
80
60
40
20
0
Control AAV
Kir4.1 AAV
Infected neurons (%)
D
Kir4.1 normalized membrane fraction
0
1
2
3
4
*
Control AAV
Kir4.1 AAV
kDa
Control AAV
Kir4.1 AAV
230
180
E
Merge
GFP
Hoechst
1 mm
500 µm
ROSTRO
CAUDAL

### Figure S6

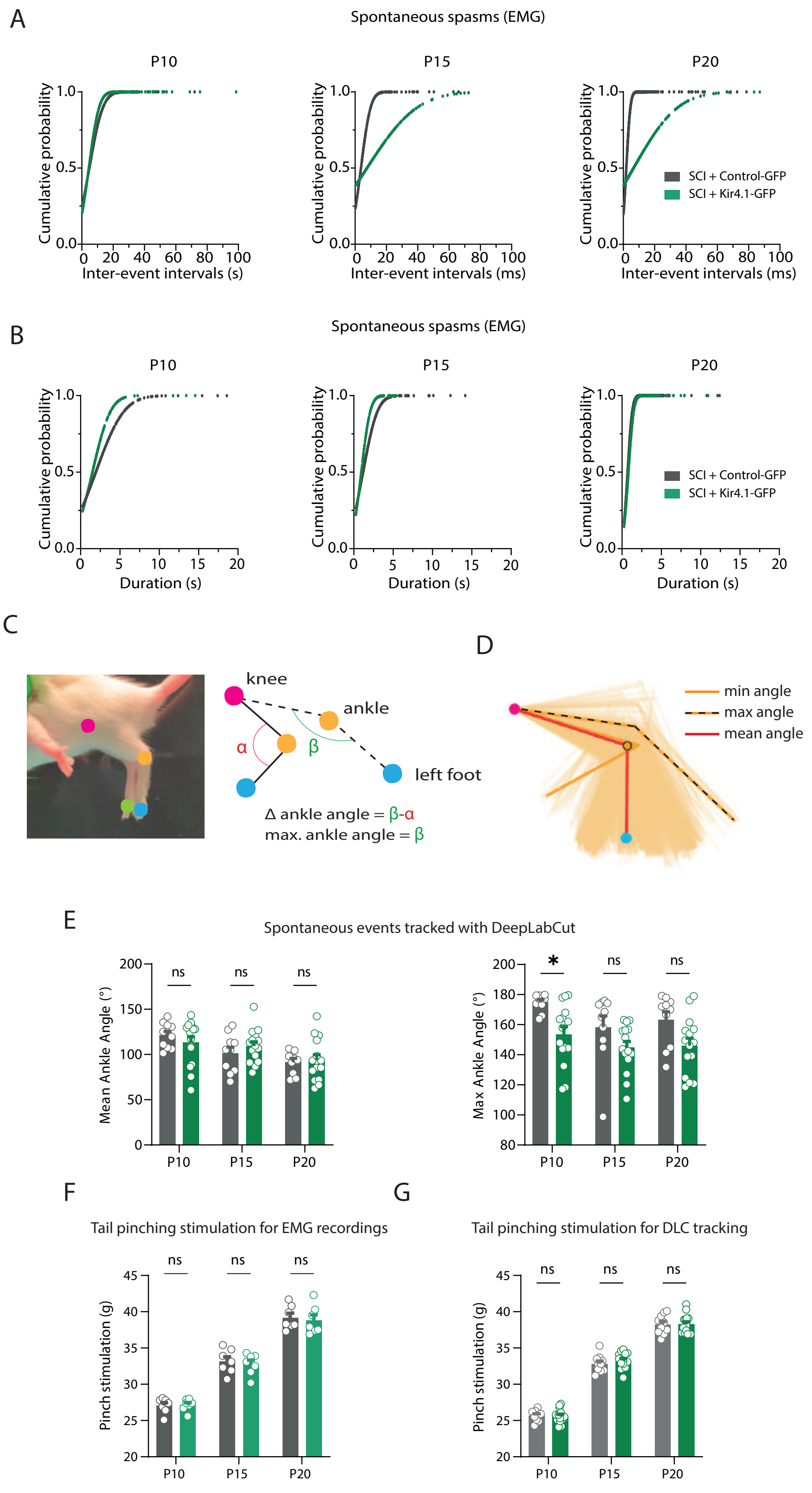

A
Spontaneous spasms (EMG)
P10
P15
P20
Cumulative probability
Inter-event intervals (s)
Inter-event intervals (ms)
SCI + Control-GFP
SCI + Kir4.1-GFP
B
Spontaneous spasms (EMG)
Duration (s)
C
knee
ankle
left foot
α
β
Δ ankle angle = β-α
max. ankle angle = β
D
min angle
max angle
mean angle
E
Spontaneous events tracked with DeepLabCut
Mean Ankle Angle (°)
Max Ankle Angle (°)
ns
*
F
Tail pinching stimulation for EMG recordings
Pinch stimulation (g)
G
Tail pinching stimulation for DLC tracking

**Figure S1 : Increased GFAP staining is associated with increased inflammatory markers the lumbar segments after thoracic SCI. (A)** Confocal images showing immunoflorescent staining of GFAP in the ventro-lateral region of the lumbar cord from control (CTL) (left) or spinal cord injured (SCI) (right) mice. Scale bar = 100 µm. GM : grey matter ; WM : white matter. **(B)** Quantification of GFAP immunoreactivity (IR) within the ventro-lateral part of the spinal cord (CTL: n = 4 mice ; SCI: n = 4 mice). **(C)** Confocal images of the MNs pool from a CTL Aldh1L1-eGFP mouse (left) and an SCI Aldh1L1-eGFP mouse (right), showing endogenous eGFP (upper middle) and immunofluorescent staining for Hoechst (top), GFAP (middle), C3d (lower middle), and the merged image (bottom). Scale bar = 15 µm **(D)** *Left*, quantification of double Aldh1L1 (+) and C3d (+) astrocytes in CTL (black) and SCI (red) mice. *Right***,** cumulative proportion of double Aldh1L1 (+) and C3d (+) astrocytes in CTL (top) and SCI (bottom) mice (CTL: n = 4 mice, n = 399 astrocytes; SCI: n = 4 mice, n = 398 astrocytes). **(E)** Confocal images of the MNs pool from a CTL Aldh1L1-eGFP mouse (left) and an SCI Aldh1L1-eGFP mouse (right), showing endogenous eGFP (upper middle) and immunofluorescent staining for Hoechst (top), STAT3 (lower middle), and the merged image (bottom). Note that the hatched white line regions of interest (ROIs) represent the nuclear segmentation (Hoechst signal) used to measure nuclear STAT3 immunoreactivity (IR). Scale bar = 10 µm. **(F)** Quantification of nuclear STAT3 IR in Aldh1L1(+) astrocytes (CTL: n = 2 mice, n = 109 astrocytes; SCI: n = 2 mice, n = 122 astrocytes). Data: mean ± S.E.M. *P < 0.05, ***P < 0.001, ****P < 0.0001 Mann-Whitney test for B, D (left) and F; Fisher's test for D (right).

**Figure S2 : Spontaneous and glutamate-evoked $Ca^{2+}$ transients in astrocytic soma and microdomains after SCI. (A)** Examples of spontaneous GCaMP6f $Ca^{2+}$ transients in astrocyte soma from CTL (top, black) and SCI (bottom, red) mice. **(B)** Percentage of active versus silent astrocytic soma from CTL (black) and SCI (red) mice (CTL : n = 54 ROIs, n = 7 mice ; SCI : n = 36 ROIs, n = 5 mice). **(C)** Quantification of the spontaneous GCaMP6f $Ca^{2+}$ transients amplitude (left), duration (middle), and frequency (right) in astrocytic soma from CTL (black) and SCI (red) mice (CTL : n = 54 ROIs, n = 7 mice ; SCI : n = 36 ROIs, n = 5 mice). **(D)** Examples of glutamate-evoked GCaMP6f $Ca^{2+}$ transients in astrocytic soma from CTL (left, black) and SCI (right, red) mice. **(E)** Quantification of the astrocytic area under the curve (AUC) of the glutamate-evoked GCaMP6f $Ca^{2+}$ transients in soma from CTL (black) and SCI (red) mice (CTL : n = 29 ROIs, n = 6 mice, SCI : n = 52 ROIs, n = 5 mice). **(F)** Examples of spontaneous GCaMP6f $Ca^{2+}$ transients in astrocytic microdomains from CTL (top, black) and SCI (bottom, red) mice. **(G)** Percentage of active versus silent astrocytic microdomains from CTL (black) and SCI (red) mice (CTL : n = 729 ROIs, n = 7 mice ; SCI : n = 788 ROIs, n = 5 mice). ROIs means regions of interest. **(H)** Quantification of the spontaneous GCaMP6f $Ca^{2+}$ transients amplitude (left), duration (middle), and frequency (right) in astrocytic microdomains from CTL (black) and SCI (red) mice (CTL : n = 729 ROIs, n = 7 mice ; SCI : n = 788 ROIs, n = 5 mice). **(I)** Examples of glutamate-evoked GCaMP6f $Ca^{2+}$ transients in astrocytic microdomains from CTL (left, black) and SCI (right, red) mice. **(J)** Quantification of the astrocytic area under the curve (AUC) of the glutamate-evoked GCaMP6f $Ca^{2+}$ transients in microdomains from CTL (black) and SCI (red) mice (CTL : n = 611 ROIs, n = 6 mice, SCI : n = 593 ROIs, n = 5 mice). ROIs mean regions of interest. All data are expressed as mean ± S.E.M.

ns: not significant, **P < 0.01, ***P < 0.001, Mann-Whitney test for C, E, H and J; Fisher's test for B and G.

**Figure S3 : Effect of propionic acid on astrocytic intracellular pH monitored with the pH-sensitive probe BCECF**. **(A)** Heatmap showing BCECF fluorescent changes in tdTomato (+) astrocytes before (left from the orange vertical dashed line) and during (right from the orange vertical dashed line) exposure of propionic acid. **(B)** Examples of BCECF signal changes in response to propionic acid exposure in astrocytes. Note, astrocytes exhibit only one response pattern : acidification (decrease in fluorescence) in response to propionic acid exposure. **(C)** Percentage of alkalynizing and acidifying responses in astrocytes from CTL mice in response to propionic acid perfusion (CTL : n = 263 ROIs, n = 4 mice). All data are expressed as mean ± S.E.M.

**Figure S4 : Blockade of Kir4.1 channels affects astrocytic $K^+$ uptake**. **(A)** Diagram illustrating the experimental procedure for recording $K^+$-inward current in astrocytes in presence of the voltage-gated sodium channel blocker, tretodotoxin (TTX). **(B)** Typical $K^+$-inward current traces under TTX following a puff application of $K^+$ (12mM) in astrocytes clamped at resting membrane potential from CTL mice in the absence (black) or presence (cyan) of the Kir4.1 channels blocker, Barium ($Ba^{2+}$). **(C)** Quantification of the astrocytic current peak amplitude and decay time time mediated by the $K^+$ puff in TTX (black) and TTX + $Ba^{2+}$ (cyan) (n = 8 astrocytes, n = 3 mice). All data are expressed as mean ± S.E.M. ns : not significant, ***P* < 0.01, Wilcoxon test for C.

**Figure S5 : Efficiency, specificity and diffusion of the viral strategy for boosting astrocytic Kir4.1. (A)** Confocal images showing immunoflorescent staining with Hoechst (left), NeuN (middle left), the endogenous GFP (middle right) and the merged signal (right) in the ventro-lateral region of a lumbar slice from a wild-type mouse (P21) injected at birth with AAV2/9-gfaABC1D-Kir4.1-GFP. The images, presented at different magnifications (top to bottom), depict the ventral section with grey matter (GM) and white matter (WM) (top), and an intermediate view of the motoneuron pool (bottom). (**B)** Confocal images showing immunoflorescent staining with Hoechst (left), GFAP (middle left), the endogenous GFP (middle right) and the merged signal (right) in the ventro-lateral region of a lumbar slice from a wild-type mouse (P21) injected at birth with AAV2/9-gfaABC1D-Kir4.1-GFP. **(C)** Proportion of GFAP (+) astrocytes (left y axis) or NeuN (+) neurons (right y axis) expressing GFP at P21 in response to birth injection of AAV-Control-GFP (grey) or AAV-Kir4.1-GFP (green). **(D)** Top : Group mean quantification of the Kir4.1 band (~ 180 kDa) normalized to the CTL group (CTL : n = 4 mice ; SCI: n = 7 mice). Kir4.1 bands were normalized to their own total protein content. Bottom : Kir4.1 pseudo-gel images from capillary western blot of lumbar segments from wild-type mice injected with Control-AAV (left) and Kir4.1-AAV (right). **(E)** Representative confocal images display Hoechst dye (left) and eGFP fluorescence signal (middle) across various coronal sections of the brain, hindbrain, and brainstem in a P21 mouse injected at birth with AAV-Control-GFP. The merged signals are shown on top. AAV2/9 with a self-complementary genome expressing eGFP was administered intrathecally (n = 2 Control-GFP mice and n = 2 Kir4.1-GFP mice). None of the four injected mice exhibited GFP fluorescence in brain structures associated with motor control. Data are presented as mean ± S.E.M. *P < 0.05, Mann-Whitney test for D.

**Figure S6 : Complementary metrics for assessing spasticity using EMG recordings and DLC tracking. (A-B)** Cumulative probability distribution for spontaneous inter-event intervals (A) and event durations (B) at P10, P15 and P20 in

SCI mice injected with control-AAV (grey) or Kir4.1-AAV (green) (SCI control AAV n=7 mice ; SCI Kir4.1 AVV n=7 mice). **(C)** Video frame of a mouse with automated body part labeling using DLC. **(D)** Representative stick diagram of hindlimb motion following tail pinch showing an example of the mean and the maximum ankle angle (β). Each stick represents a frame extrated from a video recording. **(E)** Quantification of the variation in the mean (left) and the maximum (right) ankle angle induced by tail-pinch stimulation at P10, P15 and P20 in SCI mice injected with control-AAV (grey) or Kir4.1-AAV (green). SCI + control-GFP: n = 10 mice ; SCI + Kir4.1-GFP : n=15 mice. **(F)** Quantification of the tail pinching stimulation intensity for EMG procedure (left) and for DLC acquisition (right) showing no differences between mice injected with control-AAV (grey) or Kir4.1-AAV (green) at P10, P15 and P20. SCI + control-GFP: n = 10 mice ; SCI + Kir4.1-GFP : n=15 mice. All data are expressed as mean ± S.E.M. ns : non significant, *P < 0.05, 2 way ANOVA Sidak's multiple comparaison test for E, F and G.

**Video S1 :** Typical video acquisition showing spontaneous DLC tracking on hindlimbs in a SCI mouse injected with a control-AAV.

**Video S2** : Typical video acquisition showing spontaneous DLC tracking on hindlimbs in a SCI mouse injected with a Kir4.1-AAV.
